# Whole exome sequencing identifies novel DYT1 dystonia-associated genome variants as potential disease modifiers

**DOI:** 10.1101/2020.03.15.993113

**Authors:** Chih-Fen Hu, G. W. Gant Luxton, Feng-Chin Lee, Chih-Sin Hsu, Shih-Ming Huang, Jau-Shyong Hong, San-Pin Wu

## Abstract

**Background:** DYT1 dystonia is a neurological movement disorder characterized by painful sustained muscle contractions resulting in abnormal twisting and postures. In a subset of patients, it is caused by a loss-of-function mutation (ΔE302/303; or ΔE) in the luminal ATPases associated with various cellular activities (AAA+) protein torsinA encoded by the *TOR1A* gene. The low penetrance of the ΔE mutation (∼30-40%) suggests the existence of unknown genetic modifiers of DYT1 dystonia.

**Methods:** To identify these modifiers, we performed whole exome sequencing of blood leukocyte DNA isolated from two DYT1 dystonia patients, three asymptomatic carriers of the ΔE mutation, and an unaffected adult relative.

**Results:** A total of 264 DYT1 dystonia-associated variants (DYT1 variants) were identified in 195 genes. Consistent with the emerging view of torsinA as an important regulator of the cytoskeleton, endoplasmic reticulum homeostasis, and lipid metabolism, we found DYT1 variants in genes that encode proteins implicated in these processes. Moreover, 40 DYT1 variants were detected in 32 genes associated with neuromuscular and neuropsychiatric disorders.

**Conclusion:** The DYT1 variants described in this work represent exciting new targets for future studies designed to increase our understanding of the pathophysiology and pathogenesis of DYT1 dystonia.

## Introduction

Dystonias are a heterogeneous collection of hyperkinetic neurological movement disorders that are characterized by involuntary muscle contractions resulting in abnormal repetitive movements and postures [1, 2]. Dystonias can be acquired as the result of environmental insults (i.e. central nervous system infection, toxins, and traumatic brain injury) [2, 3] as well as inherited due to genetic mutations [4]. While several causative genes are known, the mechanisms underlying their contribution to dystonia pathogenesis and/or pathophysiology remain unclear.

Early onset torsion dystonia, or DYT1 dystonia, is the common and severe inherited dystonia [5]. It is a primary torsion dystonia, as dystonia is the only clinical symptom present in patients and it is inherited in a monogenic fashion. The majority of DYT1 dystonia cases are caused by the autosomal dominantly inherited deletion of a GAG codon (c.904_906/907_909ΔGAG) from the *TOR1A* gene, which removes a glutamic acid residue (ΔE302/303; or ΔE) from the C-terminus of the encoded luminal ATPase torsinA [6, 7]. The ΔE mutation is considered a loss-of-function mutation because homozygous torsinA-knockout and homozygous torsinA^Δ^ -knockin mice both die perinatally and exhibit neurons with abnormal blebbing of the inner nuclear membrane into the perinuclear space of the nuclear E mutation impairs the ability of torsinA to interact with its major binding partners the inner nuclear membrane protein lamina-associated polypeptide 1 (LAP1) and the endoplasmic reticulum/outer nuclear membrane protein luminal domain-like LAP1 (LULL1) [9], which stimulates the ability of torsinA to hydrolyze ATP above negligible background levels *in vitro* [10].

Surprisingly, only ∼30-40% of individuals heterozygous for the ΔE develop DYT1 dystonia despite the presence of abnormalities in brain metabolism and the cerebellothalamocortical pathway in all carriers [11–15]. Collectively, these clinical findings demonstrate that the presence of the ΔE mutation results in abnormal brain function regardless of whether or not an individual develops DYT1 dystonia. Moreover, they suggest the hypothesis that the penetrance of the ΔE mutation may be influenced by additional as-of-yet unknown genetic factors.

Consistent with this hypothesis, recent research shows that genetic background modulates the phenotype of a mouse model of DYT1 dystonia [16]. In addition, expression profiling in peripheral blood harvested from human DYT1 dystonia patients harboring the ΔE mutation and asymptomatic carriers revealed a genetic signature that could correctly predict disease state [17]. The functional classification of transcripts that were differentially regulated in DYT1 dystonia patients relative to unaffected carriers identified a variety of potentially impacted biological pathways, including cell adhesion, cytoskeleton organization and biogenesis, development of the nervous system, G-protein receptor signaling, and vesicle-mediated pathway/protein transport. Since these biological pathways have all been previously associated with torsinA function [4, 18–20], we hypothesize that the penetrance of the ΔE mutation and therefore the development of DYT1 dystonia may depend upon the presence or absence of variants in genes that encode proteins that influence biological pathways associated with torsinA function. Below, we describe the use of whole exome sequencing (WES) to identify genetic variants in DYT1 dystonia patients but neither unaffected ΔE mutation carriers nor the unaffected control.

## Materials and methods

### Human Subjects

This study recruited 11 human subjects, including two patients from two separate families of Taiwanese ancestry. All subjects (or legal guardians) gave their written informed consent for participation and the study was approved by the Institutional Review Board of the Tri-Service General Hospital at the National Defense Medical Center in Taipei, Taiwan (IRB# 1-107-05-164). Detailed clinical information was obtained from corresponding clinicians and medical records.

### Purification of genomic DNA and RNA from Isolated Human Blood Leukocytes

Genomic DNA was purified from human leukocytes using the MagPurix^®^ Blood DNA Extraction Kit LV and run in the MagPurix 24^®^ Nucleic Acid Extraction System (Labgene Scietific^®^, SA, Châtel-Saint-Denis, Switzerland) following the instructions provided by the manufacturer. Total RNA was purified using Tempus™ Spin RNA Isolation Kit. Reverse transcription and cDNA synthesis were performed by QuantiTect^®^ Reverse Transcription Kit.

### Sanger Sequencing of the TOR1A gene

The DNA encoding portions of the *TOR1A* gene was PCR products amplified from the genomic DNA using the following primer pairs: 1) *TOR1A* (c.646G>C, D216H)- F: TAATTCAGGATCAGTTACAGTTGTG and –R: TGCAGGATTAGGAACCAGAT; and 2) *TOR1A* (c.904_906/907_909ΔGAG, Δ302/303E)- F: GTGTGGCATGGATAGGTGACCC and –R: GGGTGGAAGTGTGGAAGGAC. From the cDNA using the following primer pairs: Transcriptome(873bp) -F: ATCTACCCGCGTCTCTAC and –R: ATAATCTAACTTGGTGAACA; The resulting PCR products were purified using QIAquick PCR Purification Kit (Qiagen^®^) and then undergoing Sanger sequencing (Genomics^®^, Taipei, Taiwan).

### Whole exome sequencing

Purified human genomic DNA was sheared into ∼150-200 base-pair fragments using the S220 Focused-Ultrasonicator (Covaris, Woburn, Massachusetts) according to the instructions provided by the manufacturer. SureSelectXT Human All Exon V6 +UTR (Agilent Technologies, Santa Clara, CA) was then used to perform exome capture and library preparations The library were then sequenced using a NovaSeq 6000 System (Illumina, San Diego, CA) with 150 base-pair reads and output data up to 10 Gb per sample. After sequencing, Genome Analysis Toolkit (GATK) best practices workflows of germline short variant discovery (https://software.broadinstitute.org/gatk) was used to perform variant calling with default parameters [21]. Briefly, the Burrows-Wheeler Aligner was first used to align the sequenced exomes with the most up-to-date human genome reference build (hg38) (“GRCh38 - hg38 - Genome - Assembly - NCBI”. ncbi.nlm.nih.gov). Next, duplicate reads were removed using Picard after which the GATK was used to perform local realignment of the sequenced exomes with the reference genome and base quality recalibration. Then, GATK-HaplotypeCaller was used to call germline SNPs (single-nucleotide polymorphism) and indels. After variant calling, ANNOVAR was used to variant annotation [22] with database, include refGene, clinvar_20170905 (https://www.ncbi.nlm.nih.gov/clinvar/), avsnp150, dbnsfp33a, gnomad_genome, dbscsnv11. Annotated variants were selected with the following criteria: (1) filtering with exonic region, (2) removing synonymous mutation, (3) read depth ≥20. Next, we categorized these filtered variants according to the principle of inheritance. Finally, the variants of interest were validated by manually viewing them in the Integrative Genomics Viewer. All of the whole exome sequencing data generated in this study are deposited online at GenBank (https://www.ncbi.nlm.nih.gov/sra/PRJNA523662).

## Results

### Clinical Observations

Patient 1 was a 12-year-old male of Taiwanese descent who initially presented with waddling gait at seven years of age, which progressed to upper limb tremor and pronation within a few months. Over time, the patient sequentially displayed head tilt, scoliosis, kyphosis, repetitive and active twisting of his limbs. Five years after the onset of his symptoms, the patient showed generalized and profound muscle twisting and contraction, including dysarthria and dysphagia. The patient now presents with a sustained opitoshtonous-like posture and needs full assistance with executing his daily routines. Unfortunately, the patient did not benefit greatly from medical treatment and he refused deep brain stimulation due to the risks associated with the necessary surgical procedure. Neither he nor his family had a prior history of dystonia-related neurological movement disorders. Medical records from the hospital where Patient 1 received care prior to this study indicate that the patient lacks any mutations in his *FXN* or *THAP1* loci, which are both differential diagnoses of genetic, progressive, and neurodegenerative movement disorders [23, 24].

Patient 2 was a 40-year-old male of Taiwanese descent who initially presented with mild foot dystonia followed by cervical dystonia in his early twenties. He is able to execute his daily routines as the result of medical treatment. The patient had no prior history of dystonia-related neurological movement disorders and clinical information regarding the medical history of his family is unavailable.

### Examination of Known DYT1 Dystonia-Associated Mutations

To determine if either Patient 1 (subject 1) or Patient 2 (subject 11) harbored known DYT1 dystonia-associated mutations in their genomes, we used Sanger sequencing to screen their *TOR1A* genes for the presence of the ΔE mutation. Our results show that both patients are heterozygous for the ΔE mutation (Fig 1. Two patients with their family pedigrees and Sanger sequencing data. Family pedigree (A,E) and Sanger sequencing data (B,F)). We also sequenced the *TOR1A* genes of nine other family members of Patient 1. No ΔE mutation was found in the genomes of his unaffected mother (subject 2), male sibling (subject 4) or other relatives (subject 5, 7, 8,10), while his father (subject 3), paternal aunt (subject 6), and cousin of Patient 1 (subject 9) were asymptomatic heterozygotic ΔE mutation carriers. Then, we asked if the previously described protective modifier mutation D216H was present within the *TOR1A* gene of the core family of patient 1, including subject 1,2,3,4 [25]. However, none of the family members examined were positive for the D216H mutation (Fig 1. C,D). These results suggest that Patient 1 inherited the ΔE mutation from the paternal side of his family and that the absence of DYT1 dystonia in his father, paternal aunt, and cousin cannot be attributed to the presence of the protective D216H mutation.

**Figure 1.**
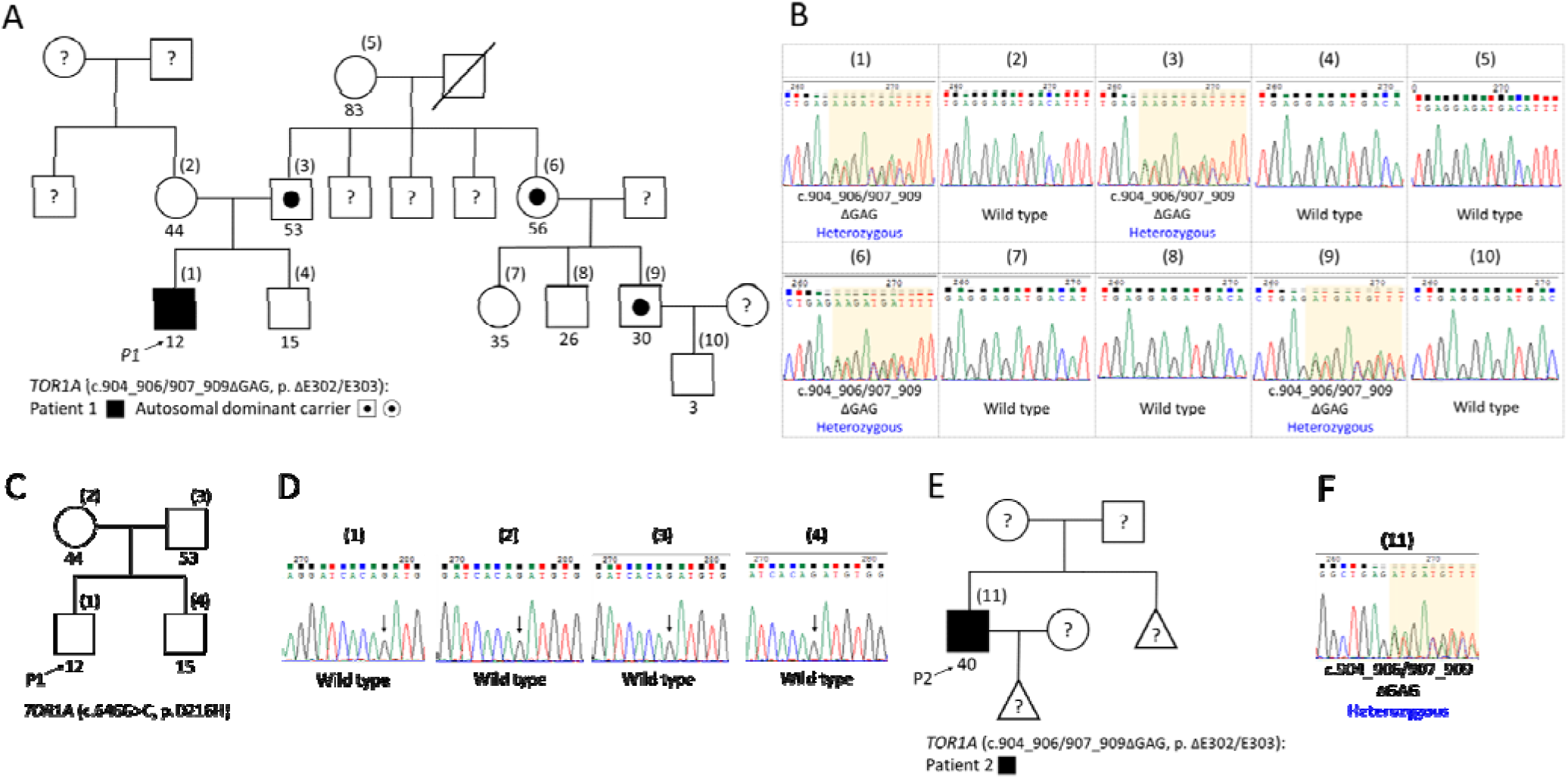
Two patients with their family pedigrees and Sanger sequencing data. Family pedigree (A,C,E) and Sanger sequencing data (B,D,F) of (A,B) 10 family members of TOR1A gene (c.904_906/907_909ΔGAG, p. ΔE302/E303), (C,D) 4 family (core family of the first family) members of TOR1A gene (c.646G>C, p.D216H) and (E,F) second patient of TOR1A gene (c.904_906/907_909ΔGAG, p. ΔE302/E303). (A,C,E) The arrows point out the two probands. The numbers within parentheses are the order of Sanger sequencing data and the numbers under the box/circle show the age (years old). The question marks within the box/circle indicate the unknown status because we don’t have the DNAs sample for study. Triangles denote lack of gender information.

### Examination of potential misregulation of alternative splicing by mutations that affect splicing signals/machinery

In addition to investigate the common genomic DNA mutation in *TOR1A* gene, mRNA extracted from leukocytes of the first patient and his parents were measured to evaluate any aberrant alternative splicing which may cause disease status [26]. There are five splicing variants and two protein production according the reference from Ensembl (human (GRCh38).p13). The results demonstrated that mRNA of *TOR1A* gene not only revealed similar gene expression level among the patient and his parents, but also showed the same GAG deletion between the patient and his father (Fig S1).

### Identification of DYT1 Dystonia-Associated Genome Variants in the *TOR1A* gene

To begin to identify potential genetic modifiers of the penetrance of the ΔE mutation, we performed WES on genomic DNA purified from blood leukocytes isolated from Patient 1 and Patient 2 as well as three asymptomatic ΔE mutation carriers from the first family (i.e. the father, paternal aunt, and cousin) and the mother of patient 1 who did not harbor the ΔE mutation in her genome. Consistent with the Sanger sequencing results described above, WES confirmed the presence of a single copy of the ΔE mutation in the exomes of Patient 1, his father, paternal aunt, and cousin as well as Patient 2, while demonstrating its absence from the mother of Patient 1 (Table 1). In addition, WES demonstrated that the D216H mutation was absent from all six exomes examined. Interestingly, three additional previously reported *TOR1A* variants, rs13300897, rs2296793 and rs1182 [27, 28], were found in the exomes of two of the asymptomatic ΔE mutation carriers and the mother from the family of Patient 1 (Table 1). However, the absence of these variants from the exomes of either Patient 1 or Patient 2 diminishes the likelihood that they are genetic modifiers of the penetrance of the ΔE mutation. Collectively, these findings motivated us to search for genome variants outside of the *TOR1A* gene that might influence the penetrance of the ΔE mutation.

**Table 1.** Genomic Variants in the Exons, Promoter Regions and 3’-UTR of the TOR1A Gene between the Patients and the Other Family Members.

| Subject Number<sup>†</sup> | Human Subject<sup>†</sup> | Variants in *TOR1A* | Site of Variant | Exonic Function | DNA Change | AA<sup>‡</sup> Change | AF<sup>‡</sup> in Person | AF in Taiwan<sup>§</sup> | AF in Dataset<sup>§</sup> | Interpretation |
| --- | --- | --- | --- | --- | --- | --- | --- | --- | --- | --- |
| (1) | Patient | rs80358233 | Exon 5 | Nonframeshift deletion | 904_906/<br>907_909ΔGAG | ΔE302/<br>303 | 0.5 | Unknown | 3.23×10<sup>-5</sup> | Pathogenic (Ozelius 2016) |
| (2) | Healthy Control | rs13300897 | Promoter | - | C>T | - | 0.5 | 0.174 | 0.1683 | Polymorphism (Vulinovic 2014) |
|  |  | rs2296793 | Exon 2 | Synonymous SNV | 246G>A | A82A | 0.5 | 0.1943 | 0.2253 | Polymorphism (Vulinovic 2014) |
|  |  | rs1182 | 3'-UTR | - | C>A | - | 0.5 | 0.178 | 0.1666 | Possible modifier (Siokas 2017) |
| (3) | Asymptomatic Carrier | rs13300897 | Promoter | - | C>T | - | 0.5 | 0.174 | 0.1683 | Polymorphism (Vulinovic 2014) |
|  |  | rs2296793 | Exon 2 | Synonymous SNV | 246G>A | A82A | 0.5 | 0.1943 | 0.2253 | Polymorphism (Vulinovic 2014) |
|  |  | rs80358233 | Exon 5 | Nonframeshift deletion | 904_906/<br>907_909ΔGAG | ΔE302/<br>303 | 0.5 | Unknown | 3.23×10<sup>-5</sup> | Pathogenic (Ozelius 2016) |
|  |  | rs1182 | 3'-UTR | - | C>A | - | 0.5 | 0.178 | 0.1666 | Possible modifier (Siokas 2017) |
| (6) | Asymptomatic Carrier | rs13300897 | Promoter | - | C>T | - | 0.5 | 0.174 | 0.1683 | Polymorphism (Vulinovic 2014) |
|  |  | rs2296793 | Exon 2 | Synonymous SNV | 246G>A | A82A | 0.5 | 0.1943 | 0.2253 | Polymorphism (Vulinovic 2014) |
|  |  | rs80358233 | Exon 5 | Nonframeshift deletion | 904_906/<br>907_909ΔGAG | ΔE302/<br>303 | 0.5 | Unknown | 3.23×10<sup>-5</sup> | Pathogenic (Ozelius 2016) |
| (9) | Asymptomatic Carrier | rs80358233 | Exon 5 | Nonframeshift deletion | 904_906/<br>907_909ΔGAG | ΔE302/<br>303 | 0.5 | Unknown | 3.23×10<sup>-5</sup> | Pathogenic (Ozelius 2016) |
| (11) | Patient | rs80358233 | Exon 5 | Nonframeshift deletion | 904_906/<br>907_909ΔGAG | ΔE302/<br>303 | 0.5 | Unknown | 3.23×10<sup>-5</sup> | Pathogenic (Ozelius 2016) |

† Subject 1: first patient, subject 2: the mother of the first patient, subject 3: the father of the first patient, subject 6: the aunt of the first patient, subject 9: the first son of the aunt of the first patient, subject 11: second patient.
‡ AA: amino acid, AF: allelic frequency.
§Taiwan biobank, https://taiwanview.twbiobank.org.tw ; gnomAD (genome aggregation database), https://gnomad.broadinstitute.org.

### Identification of DYT1 dystonia-associated Genome Variants (DYT1 variants) Outside of the TOR1A Gene

We hypothesized that candidate modifiers of the penetrance of the ΔE mutation would be those genome variants that were present in the exomes of both symptomatic patients and absent from the exomes of the asymptomatic ΔE mutation carriers and mother of the Patient 1. To begin to test this hypothesis, we examined the results of our WES for genome variants that fit this criterion and identified a total of DYT1 variants 264 variants in 195 genes. Based on their respective allele frequencies (AFs), we further classified these variants into three inheritance groups: 1) Autosomal recessive (AR); 2) Autosomal dominant (AD); and 3) *De novo* (DN) mutation (Table 2). The 53 genome variants found in 43 genes classified as AR had AFs of 1 for both patients, an AF of 0.5 or 1 for the mother of Patient 1, an AF of 0.5 for the father of Patient 1, and an AF of 0 or 0.5 for the paternal aunt and cousin of Patient 1. The 201 variants found in 149 genes classified as AD had AFs of 0.5 or 1 for both patients, and AF of 0.5 or 1 for the mother of Patient 1, and an AF of 0 for the rest of the family members of Patient 1. Finally, the 10 variants in 5 genes classified as DN had AFs of 0.5 or 1 for both patients, and were not present in any of the other exomes examined.

**Table 2.**
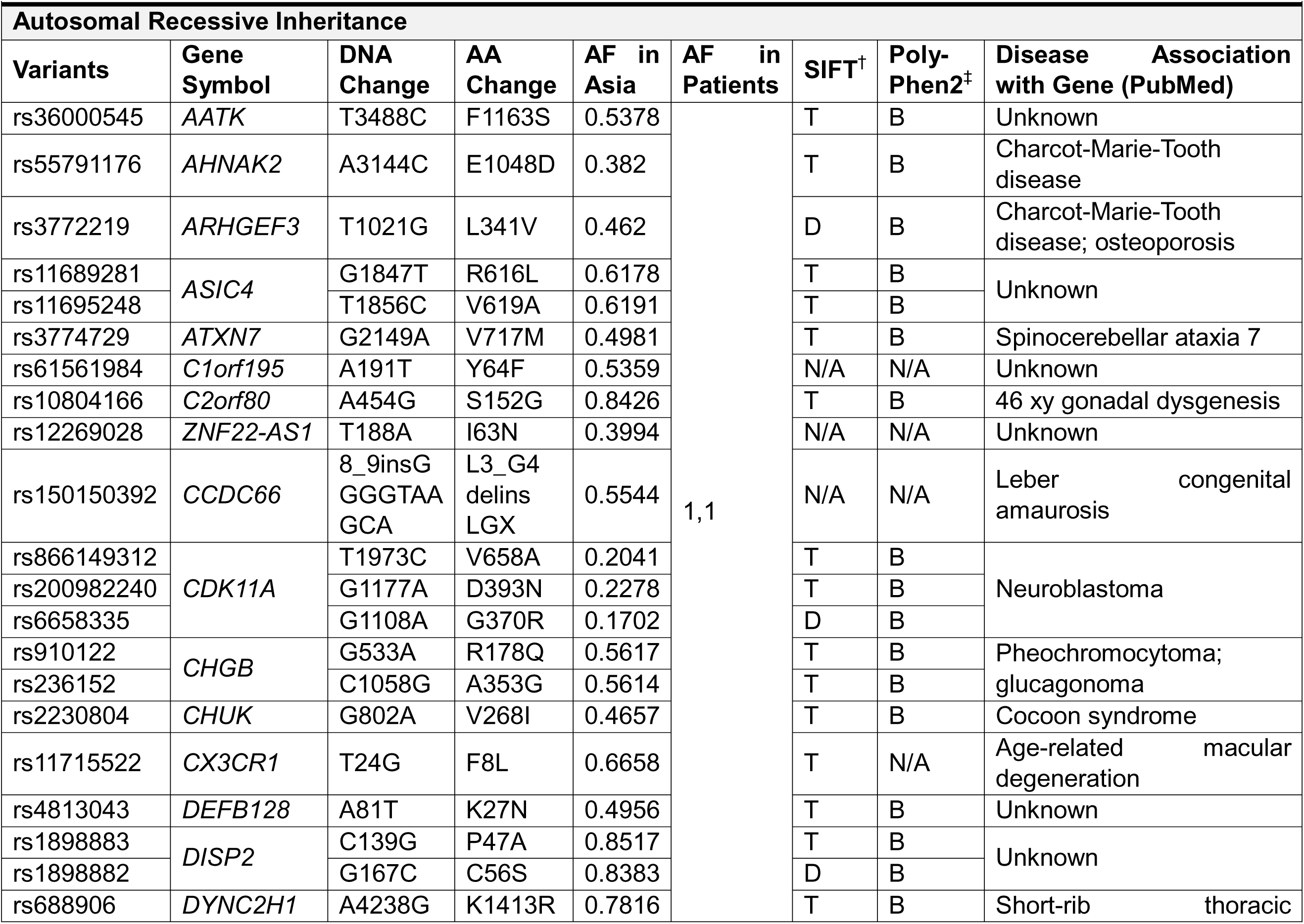
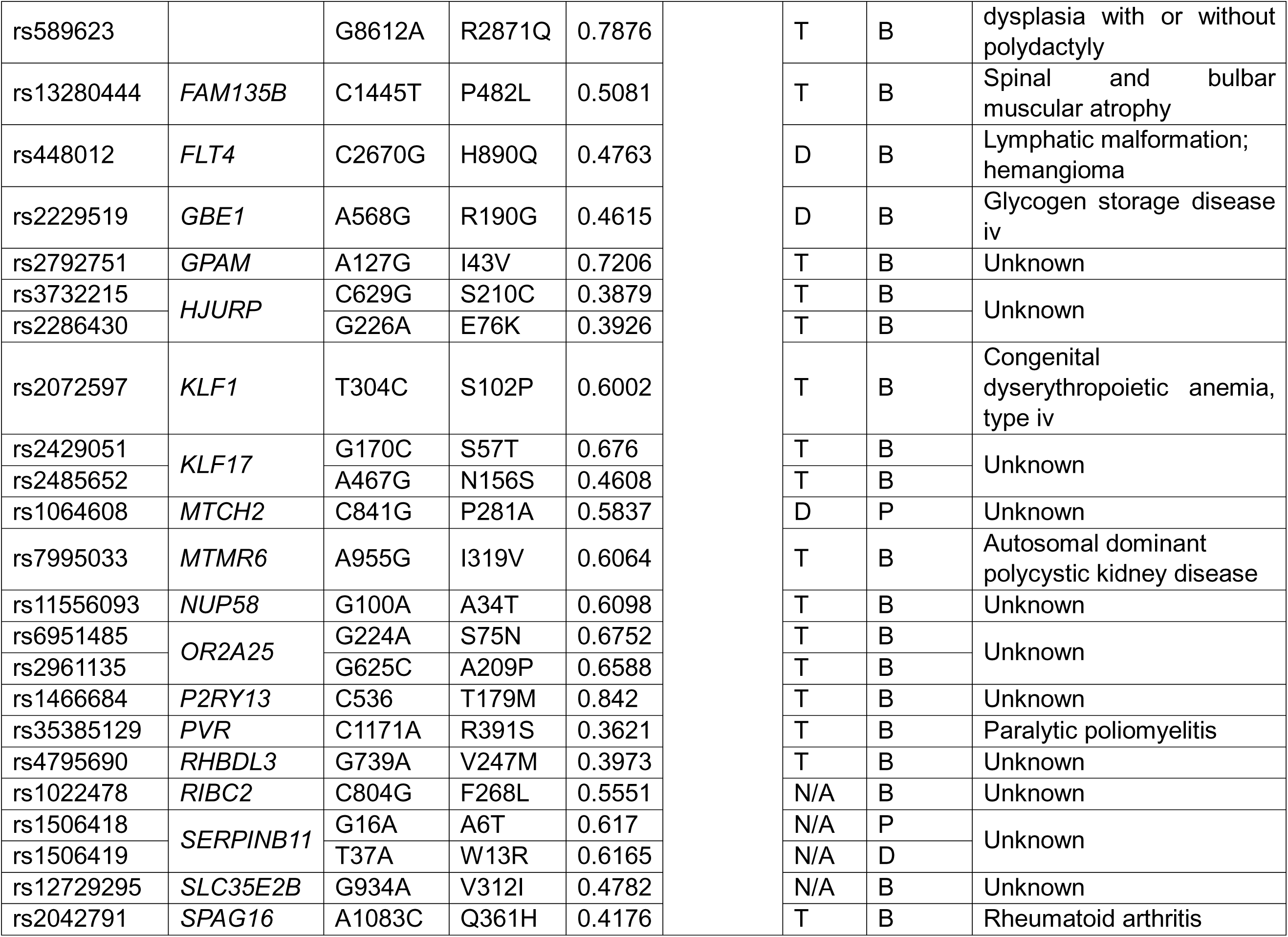
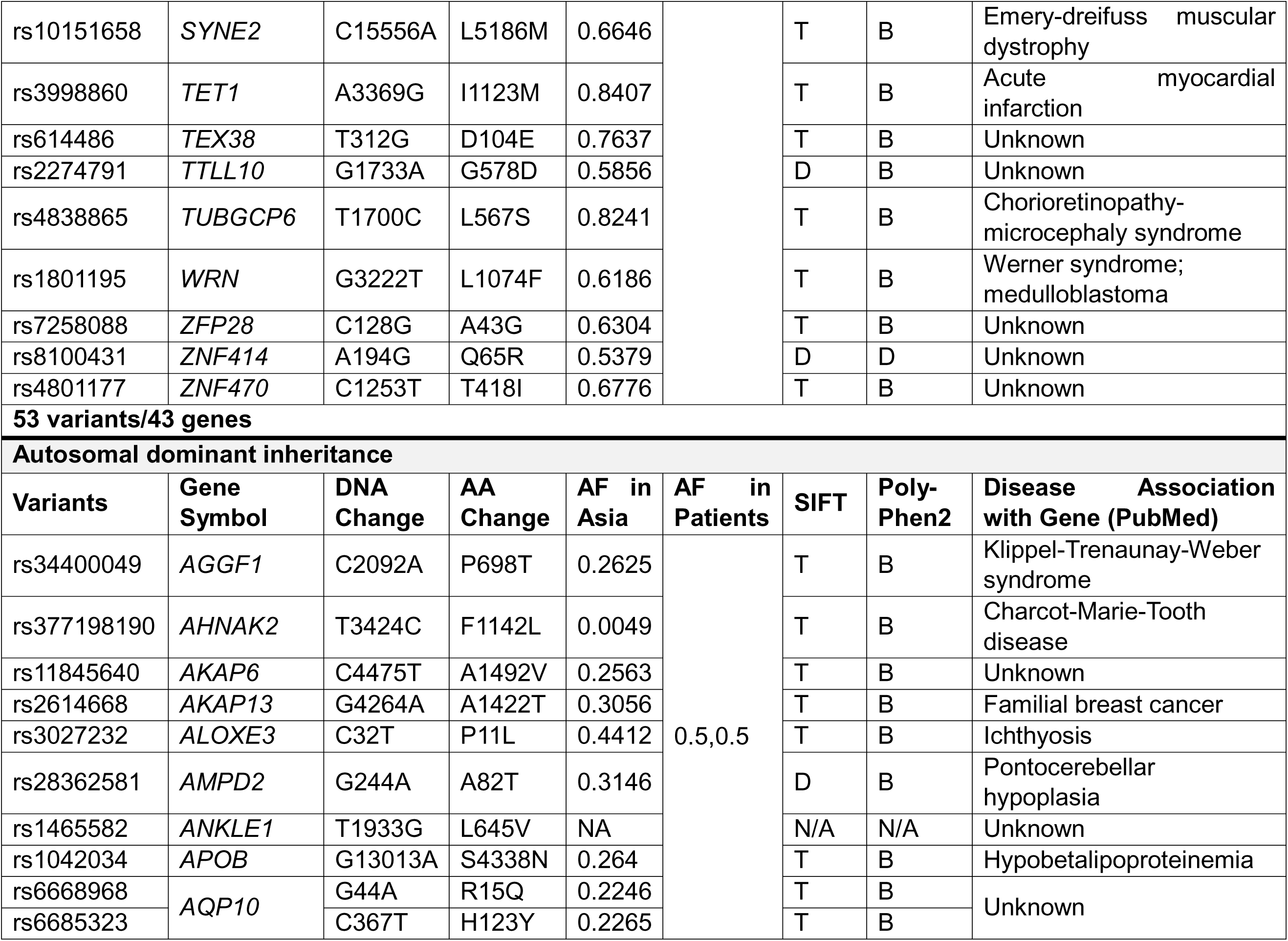
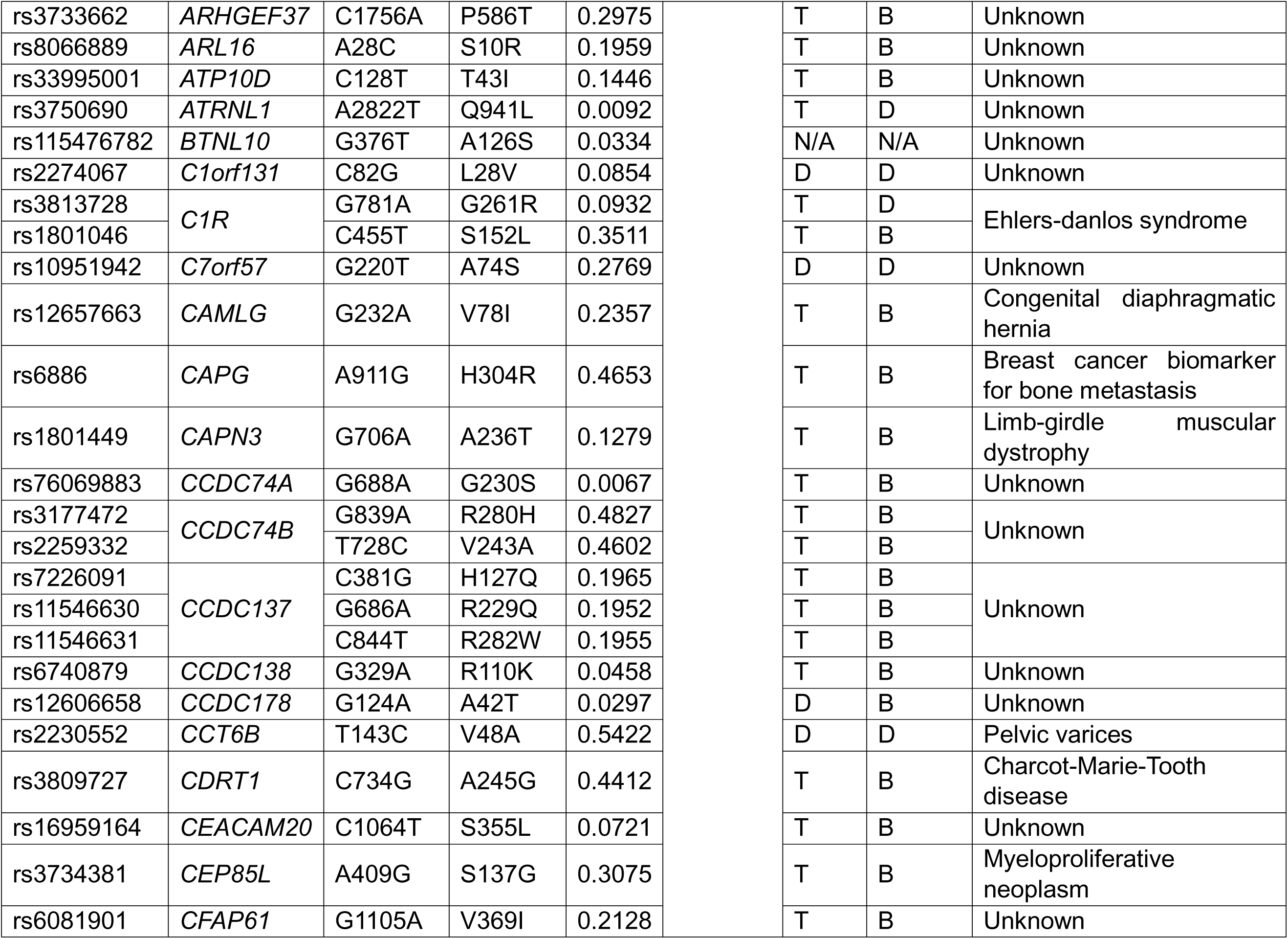
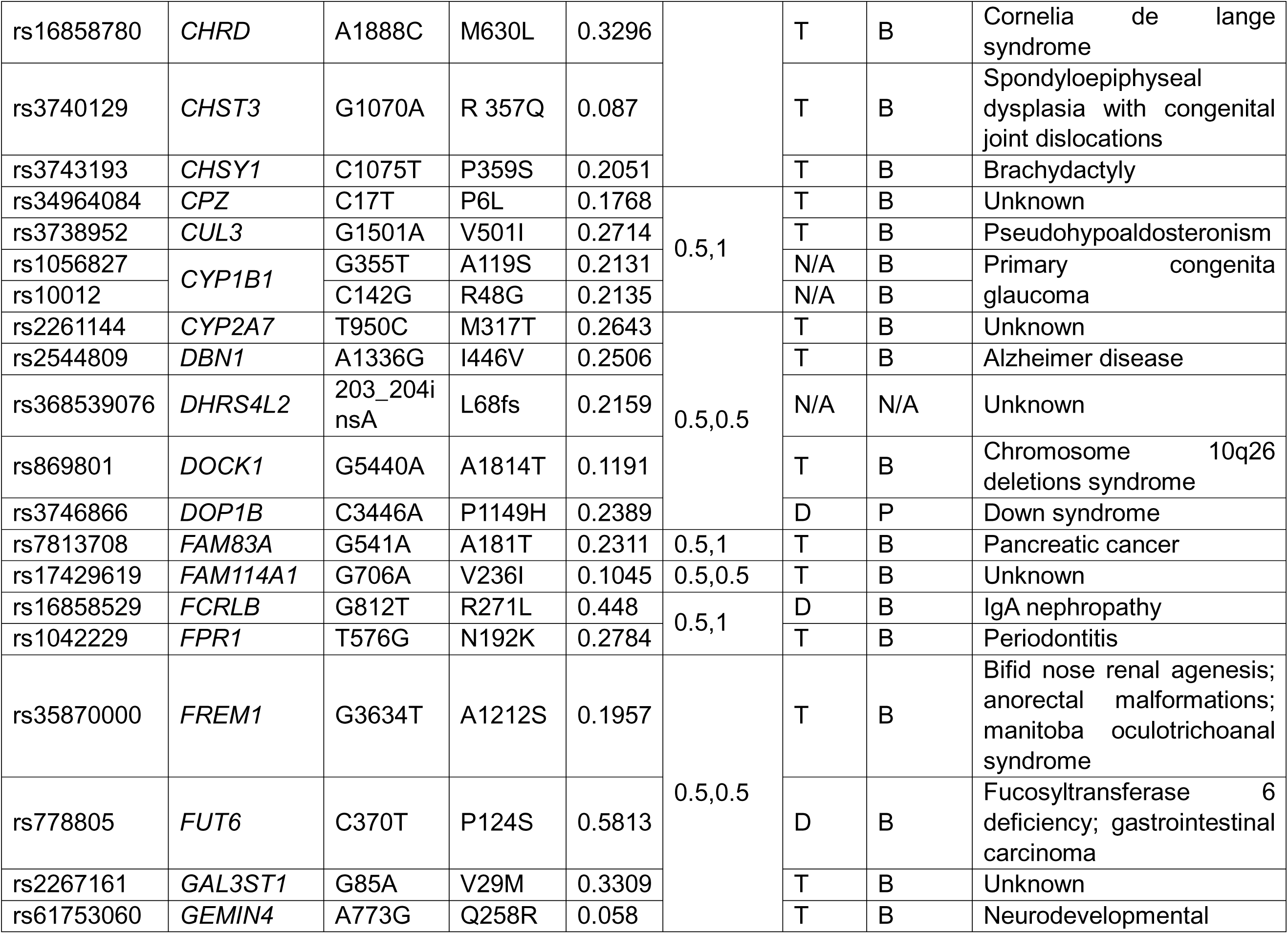
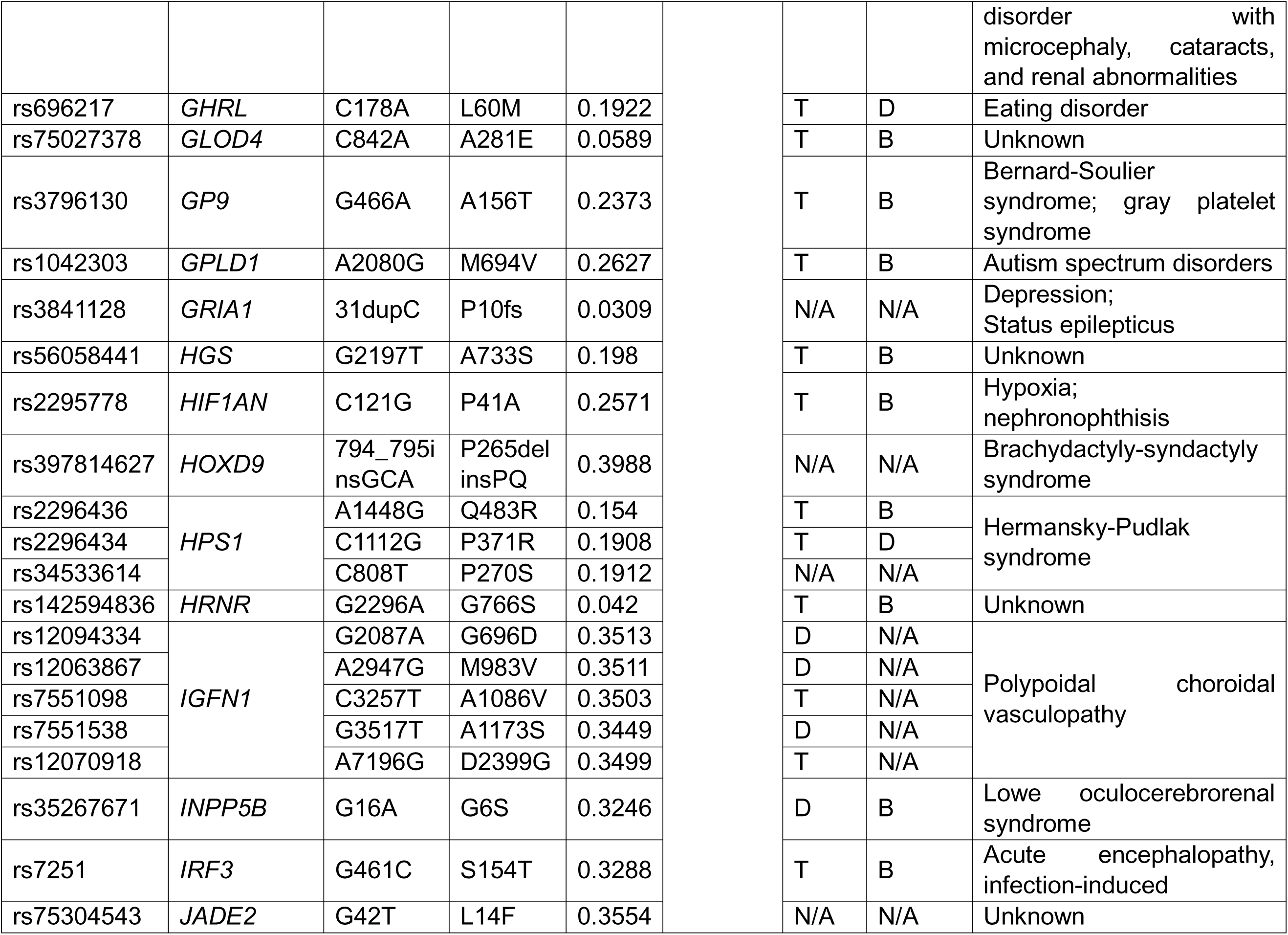
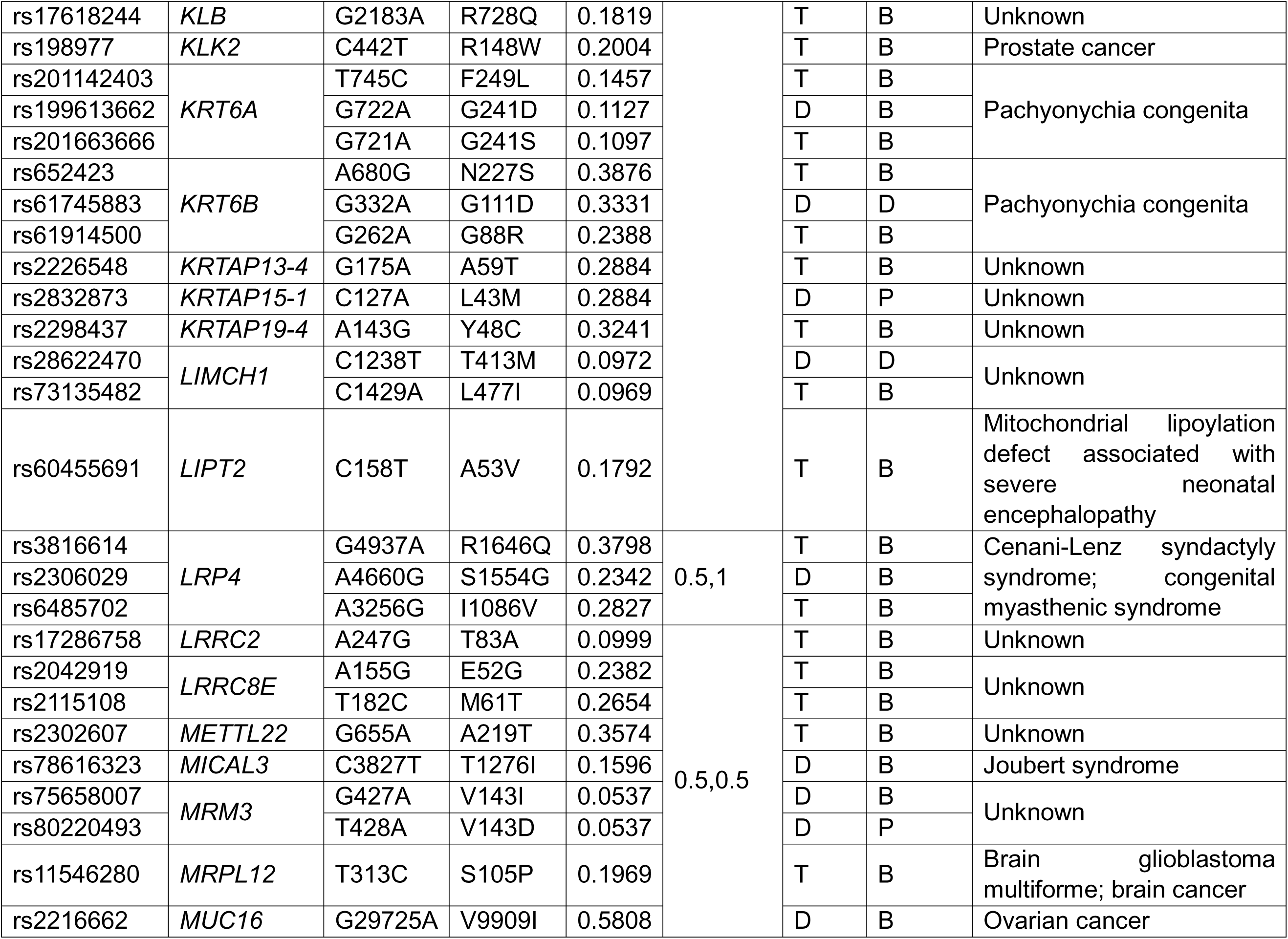
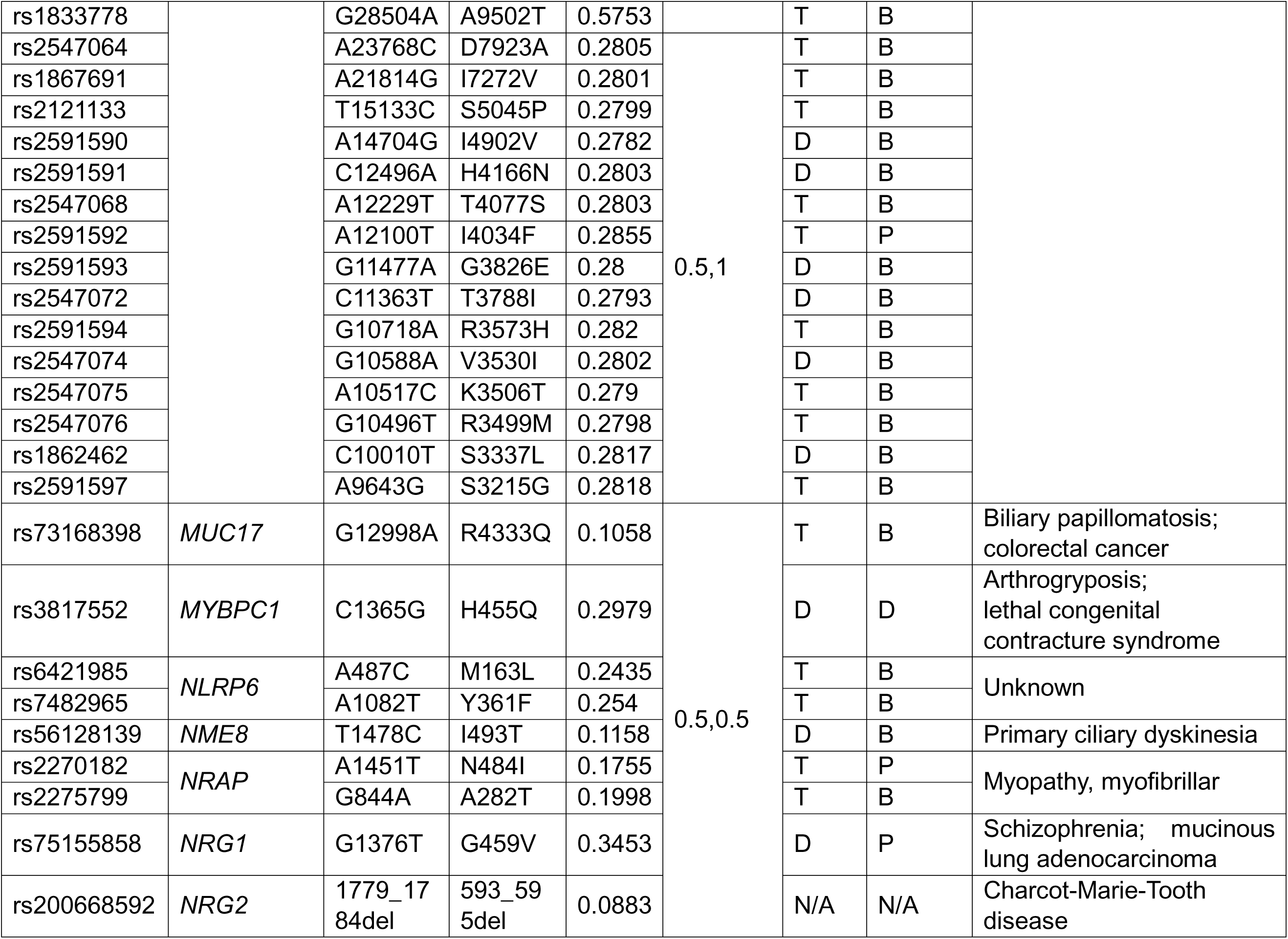
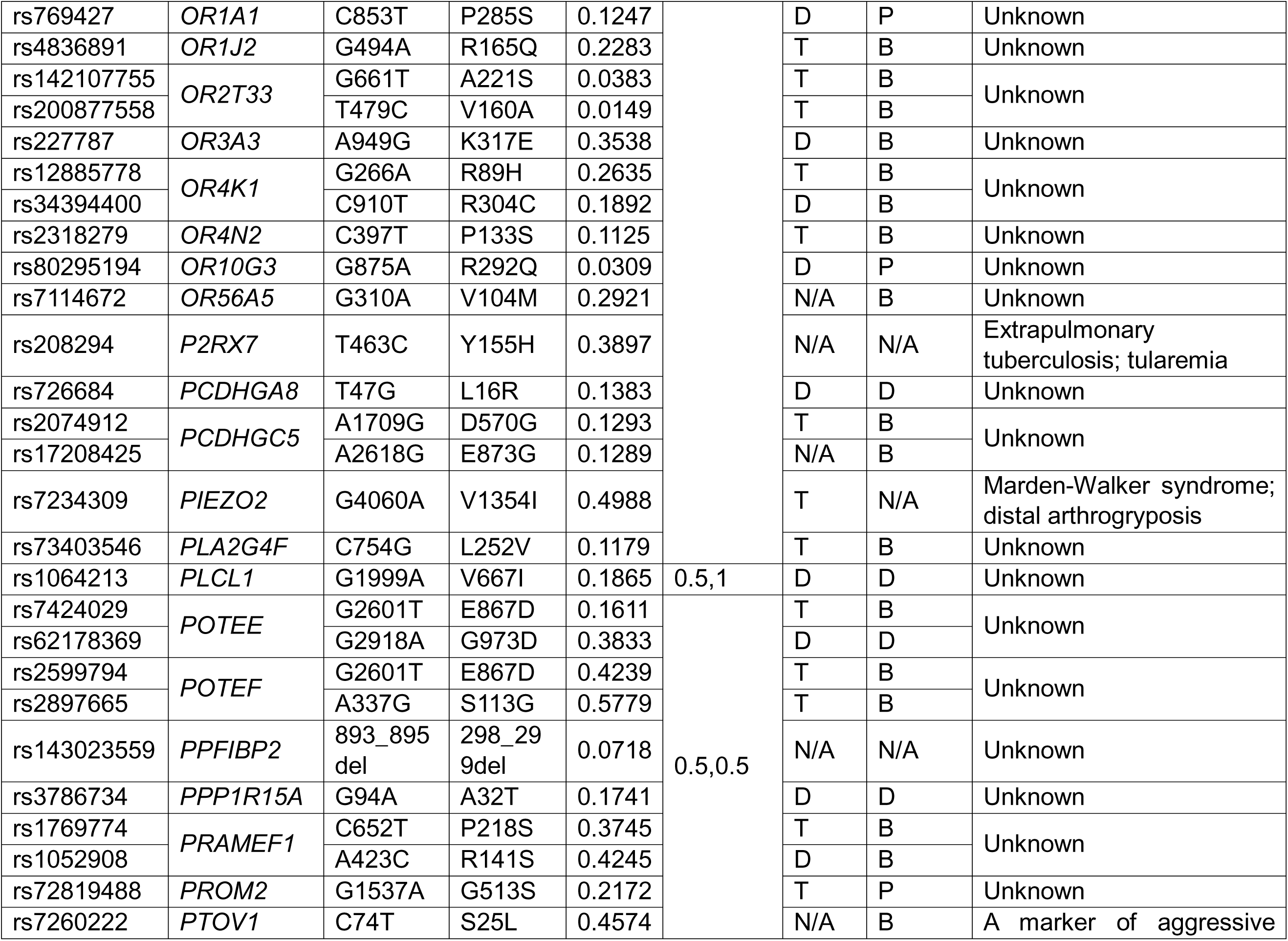
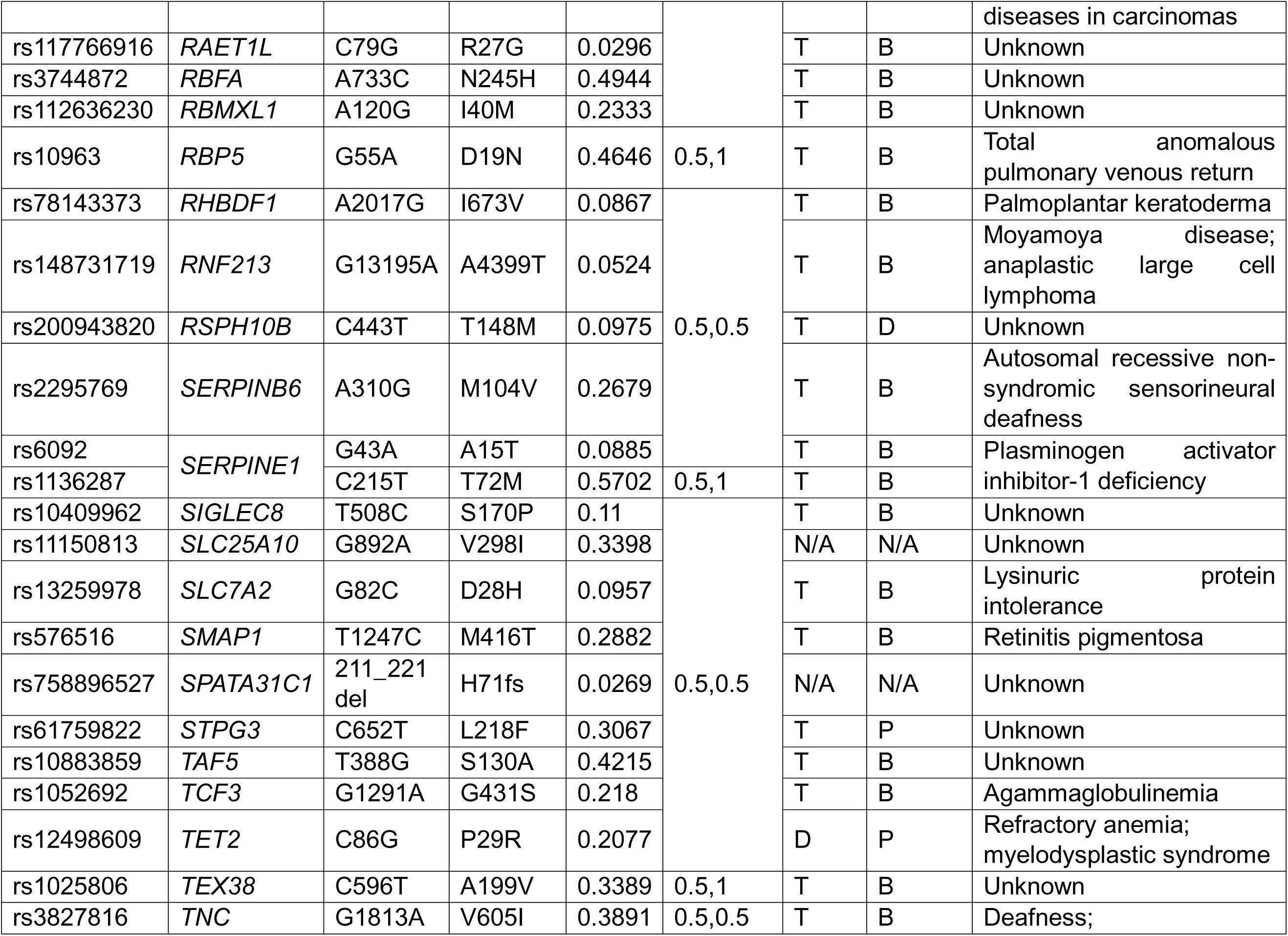
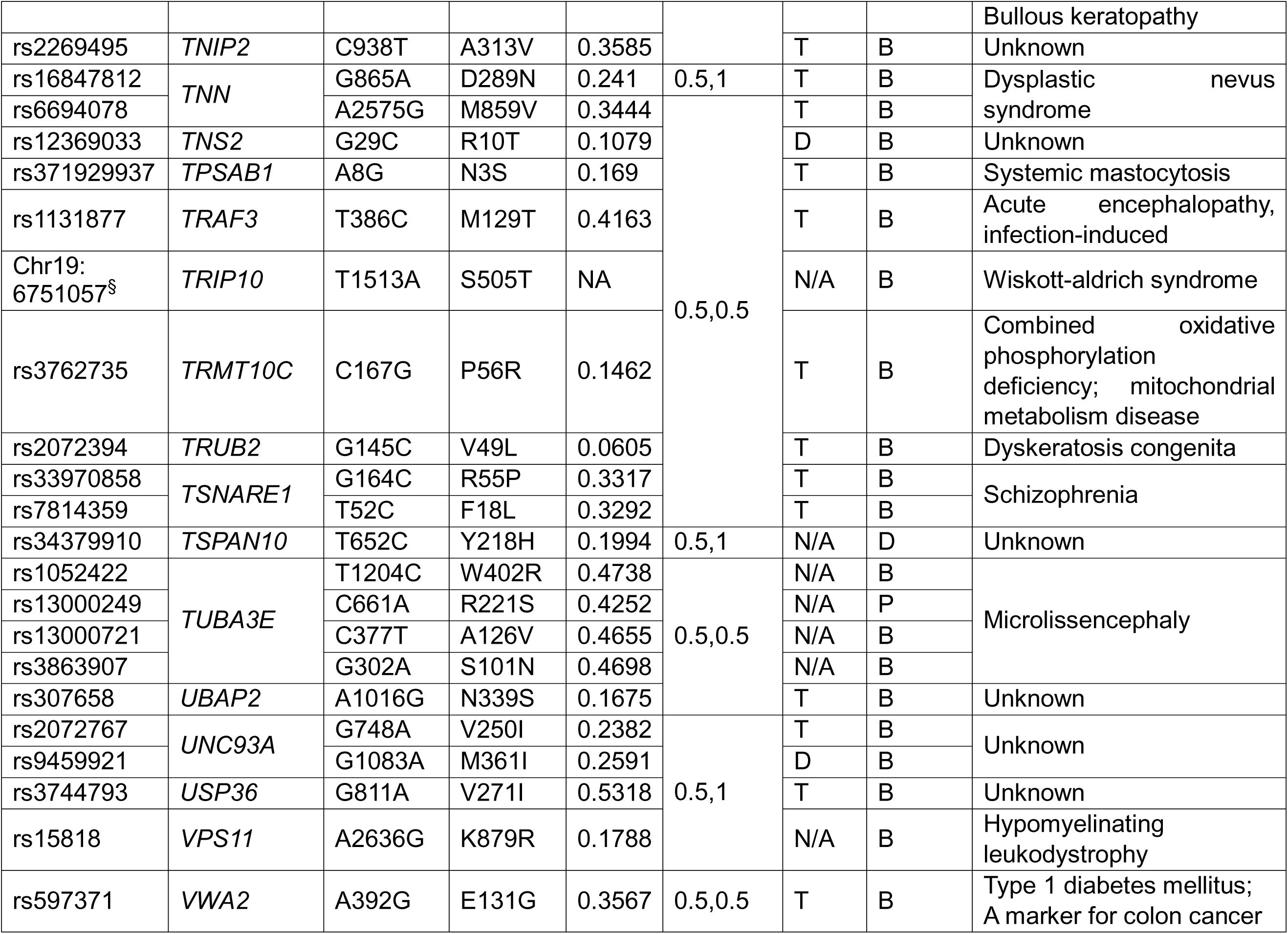
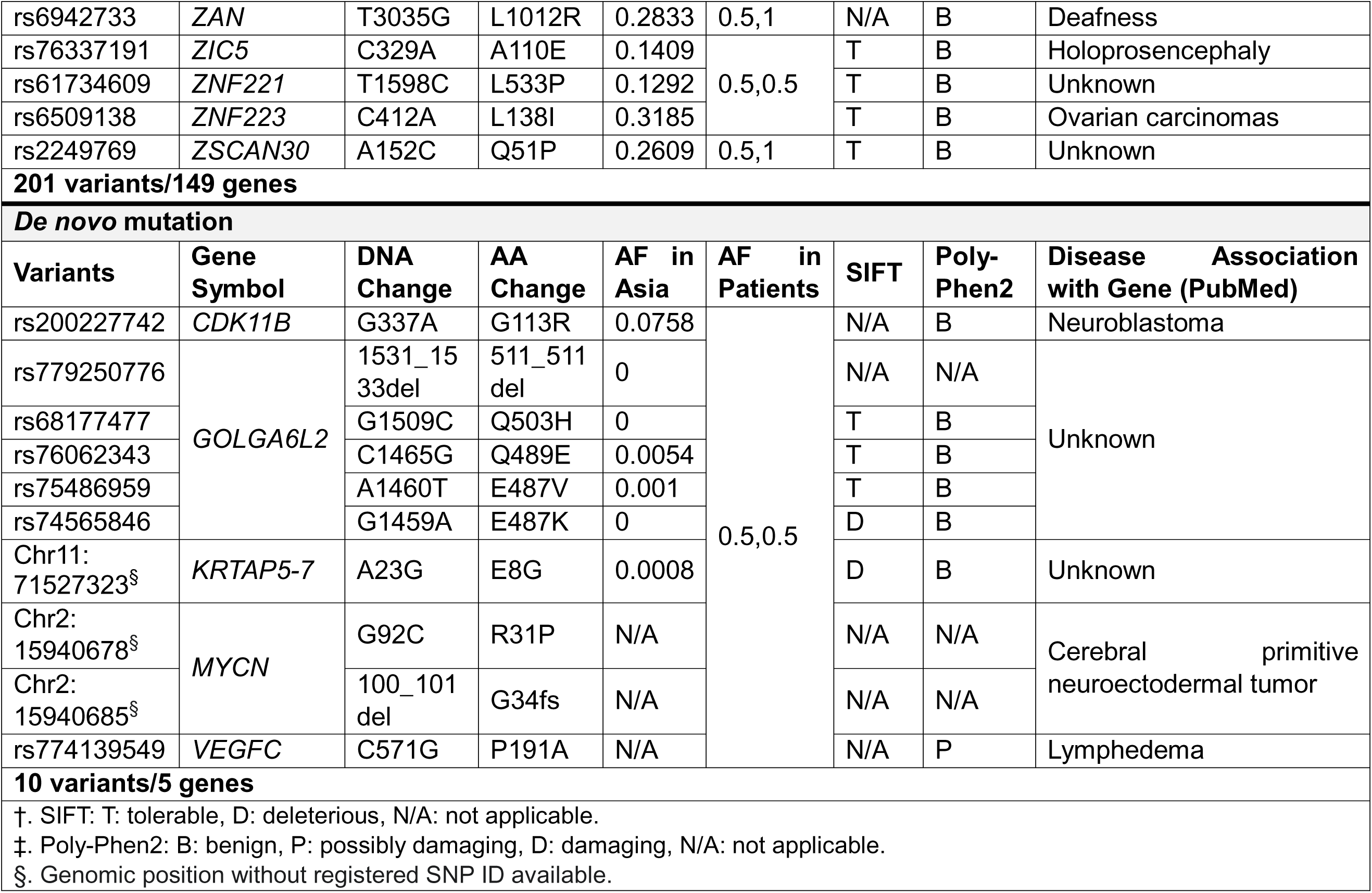
WES-identified DYT1 variants.

Of the 264 variants identified by our WES-based screen, 11 variants were predicted to be loss-of-function mutations (Table 2) by using the bioinformatics tools sorting intolerant from tolerant (SIFT) and polymorphism phenotyping (PolyPhen) [29]. Furthermore, gene ontology analysis performed using the Database for Annotation, Visualization and Integrated Discovery (DAVID) [30] identified clustered annotations of genes in which the DYT1 variants were identified by our WES-based approach. There are total 30 annotation clusters generated by this tool as listed in Table S1. Next, we filtered these categories with enrichment score>1 and *p* value<0.05 and the results were enriched for those that encode proteins that contain the epidermal growth factor-like domain (ten genes), have dioxygenase (four genes) or Rho guanyl-nucleotide exchange factor activity (four genes), or exhibit the ability to interact with the actin cytoskeleton (seven genes) (Table S1). Notably, genes for endoplasmic reticulum stress and lipid metabolism, which are linked to DYT1 functions (discussed below), also shown a trend of enrichment (Table S1). The enrichment of cytoskeleton-related genes and the known function of TOR1A in regulation of the mechanical integration of the nucleus and the cytoskeleton prompted us to look closer on the genes that harbor DYT1 variants via literature search [31–33]. There are 45 DYT1 variants in 34 genes that are associated with cytoskeleton (Table 3). In addition, 17 DYT1 variants in 16 genes are found to be linked to endoplasmic reticulum and protein and lipid metabolism (Table 4), which TOR1A is known to have functional indications at [18, 34]. Lastly, further reviewing previous studies identified 40 DYT1 variants in 32 genes that have disease associated with human neuropsychiatric disorders or neuromuscular diseases (Table 5). Taken together, our results suggest that potential regulators of the ΔE mutation may participate in the regulation of the following established cellular functions performed by torsinA: cytoskeletal organization, endoplasmic reticulum homeostasis, and protein and lipid metabolism.

**Table 3.**
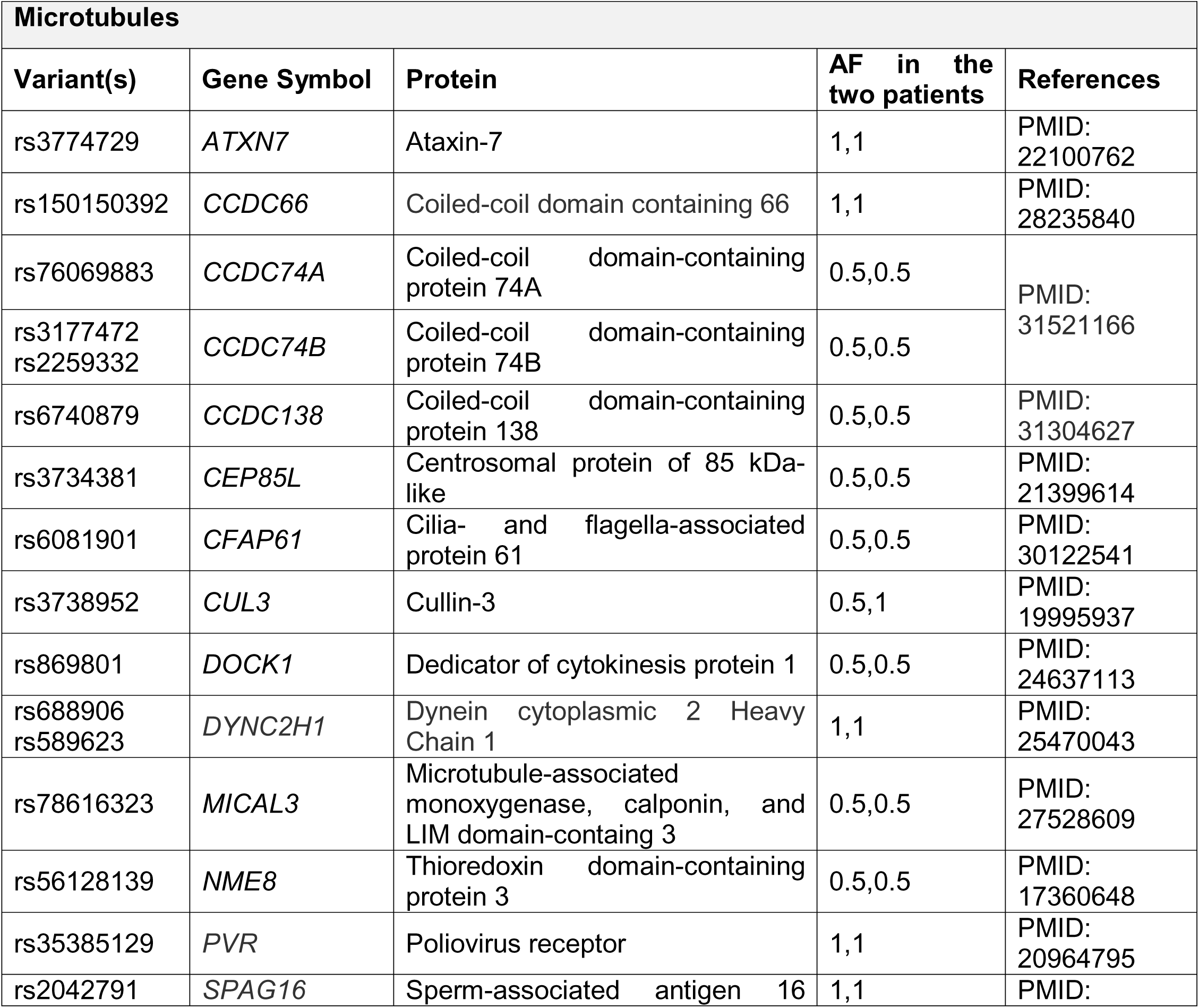
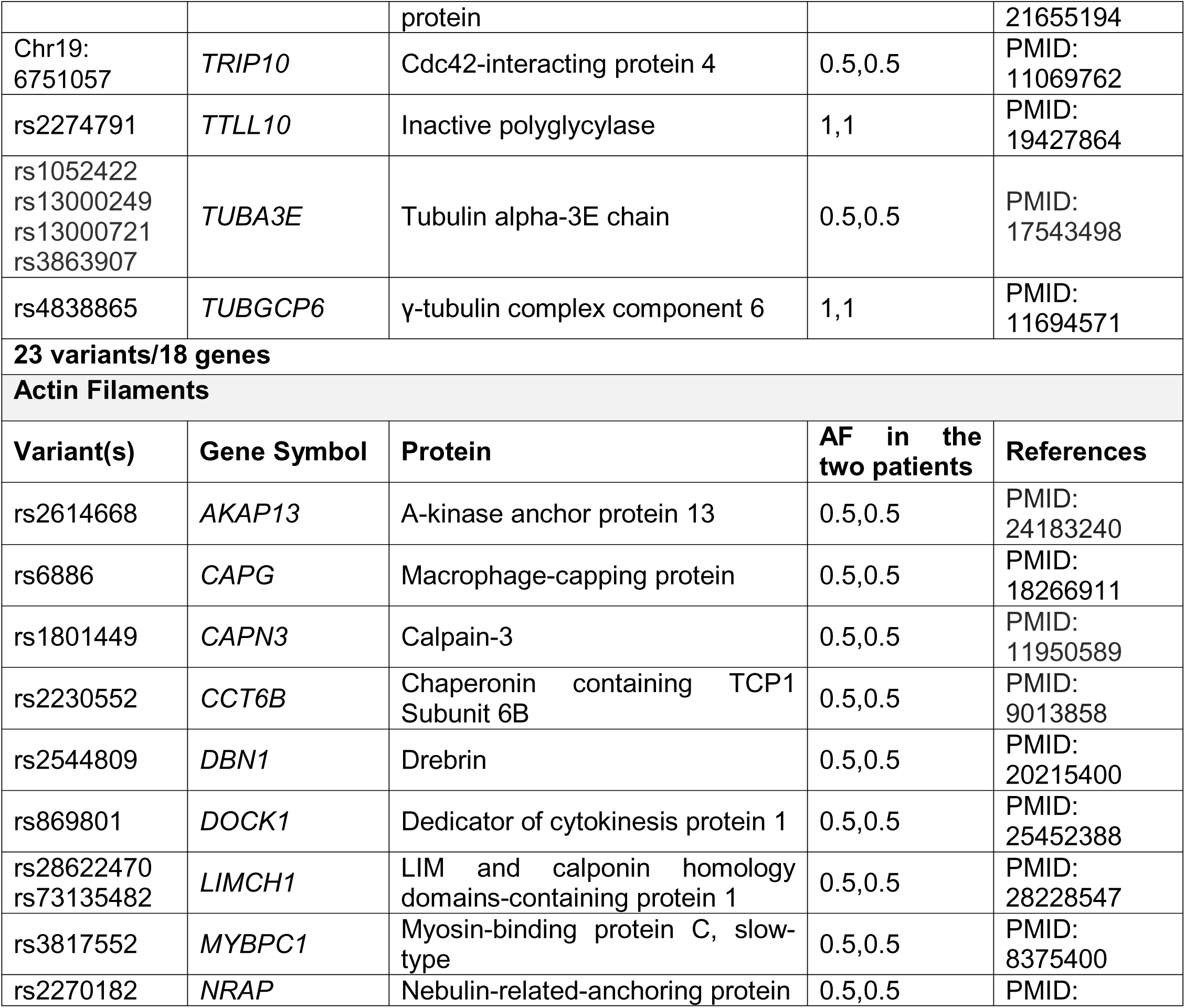
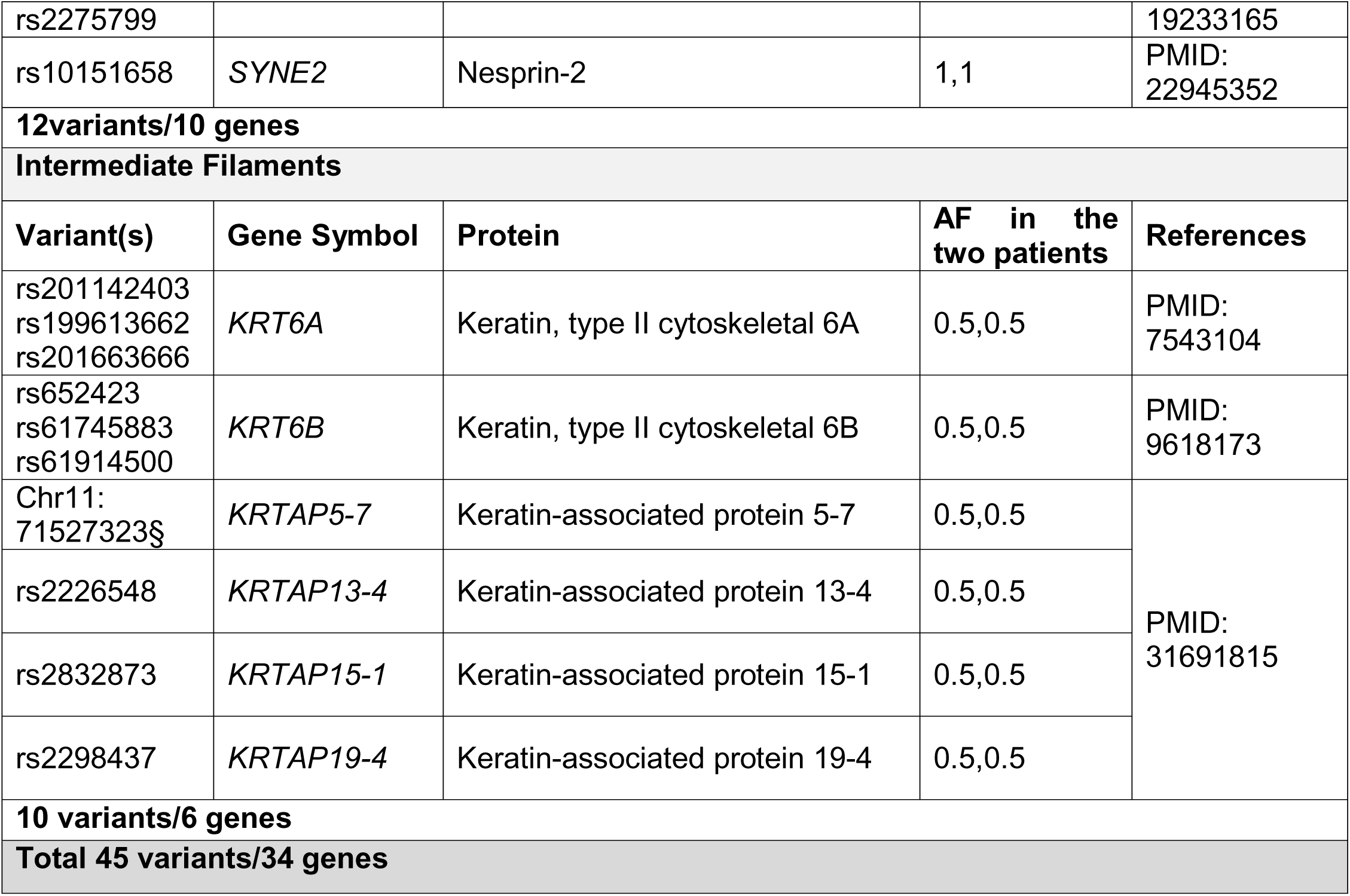
DYT1 variant-harboring genes that encode cytoskeleton-associated proteins.

**Table 4.** DYT1 variant-harboring genes that encode proteins involved in protein and lipid metabolism and endoplasmic reticulum function.

| **Endoplasmic Reticulum function and protein metabolism** |  |  |  |  |
| --- | --- | --- | --- | --- |
| **Variant(s)** | **Gene Symbol** | **Protein** | **AF in the two patients** | **References** |
| rs910122<br>rs236152 | *CHGB* | Chromogranin B | 1,1<br>1,1 | PMID:<br>31691815 |
| rs3746866 | *DOP1B* | DOP1 leucine zipper-like protein B | 0.5,0.5 | PMID:<br>16301316 |
| rs7995033 | *MTMR6* | Myotubularin-related protein 6 | 1,1 | PMID:<br>19038970 |
| rs1466684 | *P2RY13* | P2Y purinoceptor 13 | 1,1 | PMID:<br>30576484 |
| rs3786734 | *PPP1R15A* | Protein phosphatase 1 regulatory subunit 15A | 0.5,0.5 | PMID:<br>21518769 |
| **6 variants/5 genes** |  |  |  |  |
| **Lipid Metabolism** |  |  |  |  |
| **Variant(s)** | **Gene Symbol** | **Protein** | **AF in the two patients** | **References** |
| rs3027232 | *ALOXE3* | Hydroperoxide isomerase<br>ALOXE3 | 0.5,0.5 | PMID:<br>21558561 |
| rs1042034 | *APOB* | Apolipoprotein B-100 | 0.5,0.5 | PMID:<br>15797858 |
| rs1056827<br>rs10012 | *CYP1B1* | Cytochrome P450 1B1 | 0.5,1 | PMID:<br>15258110 |
| rs2261144 | *CYP2A7* | Cytochrome P450 2A7 | 0.5,0.5 | PMID:<br>21873635 |
| rs13280444 | *FAM135B* | Protein FAM135B | 1,1 | PMID:<br>21873635 |
| rs2267161 | *GAL3ST1* | Galactosylceramide sulfotransferase | 0.5,0.5 | PMID:<br>25151383 |
| rs2792751 | *GPAM* | Glycerol-3phosphate acyltransferase 1, mitochondrial | 1,1 | PMID: 18238778 |
| rs7995033 | *MTMR6* | Myotubularin-related protein 6 | 1,1 | PMID: 22647598 |
| rs73403546 | *PLA2G4F* | Cytosolic phospholipase A2 zeta | 0.5,0.5 | PMID: 21873635 |
| rs1064213 | *PLCL1* | Inactive phospholipase C-like protein 1 | 0.5,1 | PMID: 17254016 |
| **11 variants/10 genes** |  |  |  |  |

**Table 5.** DYT1 variant-harboring genes that encode proteins associated with human neuropsychiatric disorders or neuromuscular diseases

| Variant(s) | Gene Symbol | AA change | AF in Asia | Disease association | References |
| --- | --- | --- | --- | --- | --- |
| rs55791176 | *AHNAK2* | E1048D | 0.382 | Charcot-Marie-Tooth disease, Demyelinating, Type 4F | PMID: 31011849 |
| rs28362581 | *AMPD2* | A82T | 0.3146 | Pontocerebellar hypoplasia, Type 9 | PMID: 29463858 |
| rs6755527 | *ARHGEF3* | L341V | 0.462 | Charcot-Marie-Tooth disease, Type 4H | PMID: 14508709 |
| rs3774729 | *ATXN7* | V717M | 0.4981 | Spinocerebellar ataxia 7 | PMID: 30473770 |
| rs1801449 | *CAPN3* | A236T | 0.1279 | Limb-girdle Muscular dystrophy | PMID: 31540302 |
| rs200227742 | *CDK11B* | G113R | 0.0758 | Neuroblastoma | PMID: 7777541 |
| rs3809727 | *CDRT1* | A245G | 0.4412 | Charcot-Marie-Tooth disease, Type Ia | PMID: 11381029 |
| rs2544809 | *DBN1* | I446V | 0.2506 | Alzheimer disease | PMID: 28597477 |
| rs3746866 | *DOP1B* | P1149H | 0.2389 | Down syndrome | PMID: 16303751 |
| rs13280444 | *FAM135B* | P482L | 0.5081 | Spinal and bulbar muscular atrophy, X-Linked | PMID: 30391288 |
| rs2229519 | *GBE1* | R190G | 0.4615 | Polyglucosan body neuropathy, adult form | PMID: 25544507 |
| rs61753060 | *GEMIN4* | Q258R | 0.058 | Neurodevelopmental disorder with microcephaly, cataracts, renal abnormalities, and microcephaly | PMID: 25558065 |
| rs1042303 | *GPLD1* | M694V | 0.2627 | Autism spectrum disorders | PMID:<br>25448322 |
| rs3841128 | *GRIA1* | P10fs | 0.0309 | Depression;<br>Status epilepticus | PMID:<br>22057216 |
| rs7251 | *IRF3* | S154T | 0.3288 | Acute encephalopathy,<br>Infection-induced | PMID:<br>26216125 |
| rs60455691 | *LIPT2* | A53V | 0.1792 | Neonatal severe<br>encephalopathy with lactic<br>acidosis and brain<br>abnormalities and lipoic acid<br>biosynthesis defect | PMID:<br>28757203 |
| rs3816614<br>rs2300629<br>rs6485702 | *LRP4* | R1646Q<br>S1554G<br>I1086V | 0.3798<br>0.2342<br>0.2827 | Congenital myasthenic<br>syndrome | PMID:<br>28825343 |
| rs78616323 | *MICAL3* | T1276I | 0.1596 | Joubert syndrome<br>(Cerebelloparenchymal<br>disorder) | PMID:<br>26485645 |
| rs11546280 | *MRPL12* | S105P | 0.1969 | Brain glioblastoma<br>multiforme and Brain cancer | PMID:<br>26781422 |
| Chr2:<br>15940678<br>15940685 | *MYCN* | R31P<br>G34fs | NA | Cerebral primitive<br>neuroectodermal tumor | PMID:<br>28453467 |
| rs75155858 | *NRG1* | G459V | 0.3453 | Schizophrenia | PMID:<br>30500411 |
| rs2270182<br>rs2275799 | *NRAP* | N484I<br>A282T | 0.1755<br>0.1998 | Myopathy, myofibrillar | PMID:<br>30986853 |
| rs200668592 | *NRG2* | 593_59<br>5del | 0.0883 | Charcot-Marie-Tooth disease,<br>demyelinating form | PMID:<br>10369162 |
| rs148731719 | *RNF213* | A4399T | 0.0524 | Moyamoya disease | PMID:<br>29387438 |
| rs35385129 | *PVR* | R391S | 0.3621 | Paralytic poliomyelitis | PMID:<br>11597452 |
| rs10151658 | *SYNE2* | L5186M | 0.6646 | Emery-Dreifuss<br>Muscular dystrophy | PMID:<br>21496632 |
| rs1131877 | *TRAF3* | M129T | 0.4163 | Acute encephalopathy,<br>Infection-induced | PMID:<br>20832341 |
| rs33970858<br>rs7814359 | *TSNARE1* | R55P<br>F18L | 0.3317<br>0.3292 | Schizophrenia | PMID:<br>27668389 |
| rs1052422<br>rs13000249<br>rs13000721<br>rs3863907 | *TUBA3E* | W402R<br>R221S<br>A126V<br>S101N | 0.4738<br>0.4252<br>0.4655<br>0.4698 | Microlissencephaly | PMID:<br>17571022 |
| rs4838865 | *TUBGCP6* | L567S | 0.8241 | Microcephaly with<br>chorioretinopathy | PMID:<br>31077665 |
| rs15818 | *VPS11* | K879R | 0.1788 | Hypomyelinating<br>leukodystrophy | PMID:<br>27473128 |
| rs76337191 | *ZIC5* | A110E | 0.1409 | Holoprosencephaly | PMID:<br>20531442 |
| **Total 40 variants/32 genes** |  |  |  |  |  |

## Discussion

The underlying cause of phenotype variation from the same allele remains largely unknown in most cases when a particular genotype is inherited. Emerging evidence indicate that modifier genes may contribute to phenotypic variations [35]. For example, patients with thalassemia, a disorder caused by defective β-globin synthesis, have diverse clinical characteristics and variable expressivity. A number of factors underlie this phenotypic diversity, including the involvement of numerous modifier genes at other genetic loci that affect the production of β-globin [36]. Similarly, DYT1 dystonia patients have a wide spectrum of symptom severity, which reflects the incomplete penetrance of the pathogenic ΔE mutation and the variable expressivity of the disease. For most diseases, variable expressivity of the disease phenotype is the norm among individuals who carry the same disease-causing allele or alleles [37], despite the causes are not always being clear. Since the alternative splicing defects in the regulatory process may affect cellular functions and are the cause of many human diseases [26], this is not likely the potential mechanism in our cases.

In this work, we describe the identification of 264 variants in 195 genes that are associated with DYT1 dystonia. Below, we will discuss the potential implications of our results on our understanding of the pathogenesis and pathophysiology of DYT1 dystonia. Specifically, we will explore the connections between the DYT1 variants identified here and the following established cellular functions of torsinA: cytoskeletal regulation, endoplasmic reticulum stress, and lipid metabolism. In addition, we will examine the relationship revealed between DYT1 dystonia and the neuromuscular and neuropsychiatric disorders linked with the genes in which we identified DYT1 dystonia-associated genomic variants.

### DYT1 variants and the cytoskeleton

Of the 195 genes that we identified as harboring 264 DYT1 variants, 45 variants in 34 genes encode proteins that constitute or associate with the cytoskeleton (Table 3). The identification of DYT1 variants in genes encoding proteins related to cytoskeletal function is consistent with the emerging view of torsinA as a critical regulator of cellular mechanics. The first evidence to suggest that torsinA might be involved in cytoskeletal regulation was the finding that the nematode torsinA protein OOC-5 was required for the rotation of the nuclear-centrosome complex during early embryogenesis [38, 39]. In addition, the fruit fly torsinA protein torp4a/dTorsin was implicated in the regulation of the actin cytoskeleton [40]. Furthermore, the over-expression of a torsinA construct containing the ΔE mutation was shown to inhibit neurite extension in human neuroblastoma cells and to increase the density of vimentin intermediate filaments around the nucleus [41]. The relationship between torsinA and the cytoskeleton is further strengthened by reports of the impaired migration of dorsal forebrain neurons and fibroblasts from torsinA-knockout mice as well as DYT1 dystonia patient-derived fibroblasts [32, 42, 43].

More recently, torsinA was identified as a key regulator of the mechanical integration of the nucleus and the cytoskeleton via the conserved nuclear envelope-spanning linker of nucleoskeleton and cytoskeleton (LINC) complex [31–33]. The core of LINC complexes is formed by the transluminal interaction between the outer and inner nuclear membrane Klarischt/ANC-1/SYNE homology (KASH) and Sad1/UNC-84 (SUN) proteins, respectively [44]. KASH proteins interact with the cytoskeleton and signaling proteins within the cytoplasm [45], whereas SUN proteins interact with chromatin, other inner nuclear membrane proteins, and the nuclear lamina within the nucleoplasm [46].

While the precise mechanism of torsinA-mediated LINC complex regulation remains unclear, the hypothesis is supported by the finding that torsinA loss elevates LINC complex levels in the mouse brain, which impairs brain morphogenesis [47]. More recently, fibroblasts isolated from DYT1 dystonia patients were shown to have increased deformability similar to that of fibroblasts harvested from mice lacking the two major SUN proteins SUN1 and SUN2 [48].

DYT1 dystonia patient-derived fibroblasts were also shown to have increased susceptibility to damage by mechanical forces [48] strongly suggests that cellular mechanics may impact the pathogenesis and/or pathophysiology of DYT1 dystonia. All cells, including neurons, adapt their mechanical properties by converting extracellular mechanical stimuli into biochemical signals and altered gene expression through the process of mechanotransduction [49, 50]. Since mechanotransduction instructs neuronal differentiation, proliferation, and survival [51, 52], it is possible that defective mechanotransduction of neurons in the developing brain may contribute to the pathogenesis and/or pathophysiology of DYT1 dystonia. Based on the information provided above, it is intriguing that we identified DYT1 variants in the KASH protein nesprin-2-encoding *SYNE2* gene and the *NUP58* gene, which encodes the nuclear pore complex protein nup58 (Table 3 and Fig 2). In the future, it will be interesting to test if the DYT1 variants found in *SYNE2* and *NUP58* negatively impact LINC complex-dependent nuclear-cytoskeletal coupling and/or mechanotransduction.

**Figure 2.**
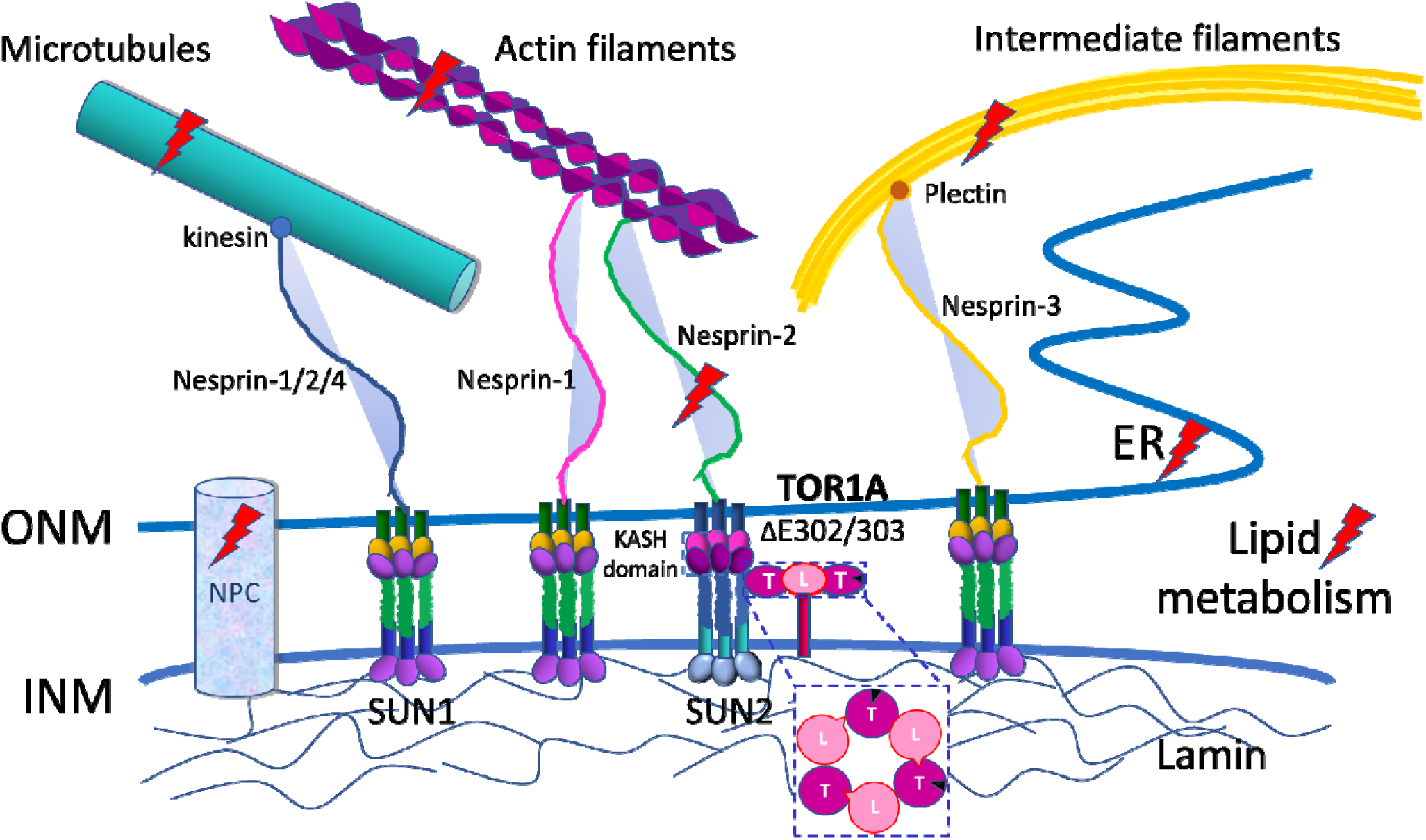
Multiple hits (variants) in the nuclear-cytoskeletal coupling network. TOR1A-LAP1 (or LULL1) heterohexamer regulates the assembly and function of LINC complex. The location of the defects at TOR1A (ΔE302/E303) and variants found in microtubules, actin filaments, intermediate filaments, nesprin-2, nuclear pore complex (NPC), endoplasmic reticulum (ER), and lipid metabolism. LINC complex (the linker of nucleoskeleton and cytoskeleton, consisting of KASH domain and SUN proteins), T (TOR1A), L (LAP1 or LULL1), KASH domain (Klarsicht, ANC-1, and Syne homology domain), SUN 1 and SUN 2 (SUN (Sad1, UNC-84) domain-containing protein 1 and 2), ONM (outer nuclear membrane), INM (inner nuclear membrane).

It is tempting to speculate that the impairment of the microtubule cytoskeleton is particularly relevant to dystonia pathogenesis given the enrichment of DYT variants that we found in genes that encode microtubule-associated proteins. Microtubules are fundamentally important for the structure and function of neurons, which are some of the most highly polarized cells in the human body [53]. Microtubules establish the polarized architecture of neurons and serve as tracks for microtubule motor proteins as they carry proteins and lipids to where they are needed for proper neuronal function. Thus, defects in microtubule dynamics and organization underly a wide array of neurological and neuropsychiatric disorders [54–56].

Consistent with our identification of 4 DYT1 variants in the *TUBA3E*, which encodes the protein α-tubulin-3E, mutations in the β-tubulin-4A-encoding *TUBB4A* gene cause another hereditary dystonia, Whispering dysphonia or DYT4 dystonia [57, 58]. These mutations result in the formation of disorganized microtubule networks and the impaired growth of neuronal processes similar to the clinical phenotypes observed in DYT4 dystonia patients [59, 60]. Future experiments designed to test the impact of the DYT1 variants in *TUBA3E* on the organization and function of neuronal microtubules will help elucidate the role of the microtubule cytoskeleton to the manifestation of DYT1 dystonia.

### DYT1 variants in association with protein synthesis and transport and ER homeostasis

Accumulating evidence indicate a role of TOR1A in the cellular protein quality control system in which TOR1A could be both substrate and effector [18]. In the 264 genome variants, we observed six variants in five genes, *CHGB, DOP1B, MTMR6, P2RY13* and *PPP1R15A*, that are annotated with protein synthesis and transport functions (Table 4). Notably, CHGB and PPP1R15A has also been linked to endoplasmic reticulum stress [61–63]. These findings support the previously proposed hypothesis that elevated levels of endoplasmic reticulum stress contributes to DYT1 dystonia pathogenesis [64–72].

TorsinA functions to protect against insults from protein aggregates in the neural system [65]. Protein aggregates are products of protein misfolding commonly seen in neurodegenerative diseases such as Alzheimer’s disease, Parkinson’s disease, amyotrophic lateral sclerosis and prion disease, which triggers endoplasmic reticulum stress response [73, 74]. In the TOR1A ΔE mutation background, we identified six candidate modifier genome variants in five genes that have known functions in endoplasmic reticulum for protein post translational modification, protein translocation and endoplasmic reticulum stress response (Table 4). Among them, DOP1B has neurological roles in both human and mice [75, 76]. Whether DOP1B’s endoplasmic reticulum cellular function has a causal effect on its neurological role remains to be investigated. Collectively, our data provide clinical indications of candidate genes and genome variants for further investigation on the underlying mechanisms of TOR1A dependent ER dysfunction in DYT1 dystonia.

### DYT1 variants and lipid metabolism

TOR1A also has a pivotal role in lipid metabolism as demonstrated by the hepatic steatosis of liver-specific torsinA-knockout mouse model [34] and the requirement for the Drosophila torsinA homologue for proper lipid metabolism in adipose tissue [77]. Because of its functional indication in lipid metabolism, TorsinA is thought to promote membrane biogenesis [19] and synaptic physiology [78]. There are 11 DYT1 dystonia associated genome variants identified in ten lipid metabolism genes *ALOXE3, APOB, CYP1B1, CYP2A7, FAM135B, GAL3ST1, GPAM, MTMR6, PLA2G4F* and *PLCL1* (Table 4), which suggest potential genetic interactions between the ΔE mutation and genome variants that might change membrane homeostasis.

TOR1A regulates lipid metabolism in both fruit flies and mammals [34, 77]. TOR1A facilitates cell growth, raises lipid content of cellular membrane and is involved in membrane expansion [77]. The linkage between the TOR1A ΔE mutation and 10 lipid metabolic genes suggest the impact on lipid metabolism associated cellular functions could be amplified by clustered mutations and genome variants. Two genes in this category have known functions in the neural system. The *GAL3ST1* gene encodes galactose-3-O-sulfotransferase 1 that involves in the synthesis of a major lipid component of the myelin sheath galactosylceramide sulfate [79]. *Gal3st1* deficient mice develop tremor, progressive ataxia, hind limb weakness, aberrant limb posture and impaired limb coordination with morphological defects in the neural system [80]. PLCL1 Involves in an inositol phospholipid-based intracellular signaling cascade. PLCL1 is phospholipase C like protein lacking the catalytic activity. PLCL1 binds and sequesters inositol triphosphates to blunt the downstream calcium signaling [81]. PLCL1 has been linked to the trafficking and turnover of GABAA receptors in neurons [82, 83]. Physiologically, loss of PLCL1 increases the incidence of chemically induced seizure in mice [84]. These findings indicate an essential role of PLCL1 in controlling the neural signaling transduction. While the functional impact of the genome variants on GAL3ST1 and PLCL1 awaits further investigation, their association with the TOR1A ΔE mutation suggests potential functional interactions between these molecules in DYT1 dystonia.

### Connections between the genes harboring DYT1 variants and their implicated neuromuscular and neuropsychiatric disorders

The loss of torsinA function in either the cerebral cortex or cerebellum result in motor dysfunction [85–87], indicating a neuronal component of TOR1A’s function in dystonia. Based on these observations, we examined the 195 genes that carry candidate ΔE mutation modifiers for their association with neuropsychiatric and neuromuscular disorders. Such link was identified in 32 genes with 40 genome variants (Table 5). These include the *AHNAK2*, *ARHGEF3*, *CDRT1*, *GBE1* and *NRG2* genes in associated with peripheral neuropathy (Charcot-Marie-Tooth disease and Polyglucosan body neuropathy, adult form). The *AMPD2*, *ATXN7* and *MICAL3* genes are linked to cerebellar diseases (Pontocerebellar Hypoplasia, type 9 and spastic paraplegia 63, autosomal recessive; Spinocerebellar ataxia 7; Joubert syndrome (cerebelloparenchymal disorder)). Lastly, the *IRF3*, *TRAF3* and *LIPT2* genes are associated with encephalopathy (acute, infection-induced; encephalopathy, neonatal severe, with lactic acidosis and brain abnormalities and lipoic acid biosynthesis defects. Overall, more than 16% of the identified 195 genes are in association with neuropsychiatric and neuromuscular diseases related disorders, demonstrating the significance of the linkage between DYT1 dystonia and these diseases.

### Study Limitations

The present study examined five individuals who have the TOR1A ΔE mutation. Among them, two have disease presentation and three are asymptomatic carriers. Furthermore, one affected patient and the three asymptomatic carriers are in the same family, which is an advantage to have a relatively close genetic background for modifier screening. Data from this family identified 1725 of genome variants as candidate modifiers. With the addition of the second affected patient, the number of candidate modifier variants were further narrowed down to 264. This number could have been reduced if data from more affected patients or asymptomatic carriers are available. Unfortunately, family members of the second affected patient declined to participate in the study. Due to the rareness of DYT1 dystonia in Taiwan, it is difficult to increase sample size within the Taiwanese population in foreseeable future. Alternatively, meta-analysis of our dataset with WES results from other populations across the world, once publicly available, may help to identify the common modifiers in the general population [88, 89].

The WES data allows identification of candidate modifiers in the coding genome. However, majority of the GWAS signals are mapped to the noncoding regions of the genome and accumulating evidence point to disease associations with the noncoding genome [90]. Mutations in the noncoding genome may impact cis-acting element functions and chromatin conformations that direct gene expression. Future inclusion of the whole genome sequencing assay may help to identify additional modifiers for the DYT1 dystonia.

## Conclusions

In summary, we propose that genome variants within nuclear-cytoskeletal coupling network may constitute potential modifier variants, which could synergistically reduce the threshold of disease onset of DYT1 dystonia and accelerates the clinical symptoms and signs of dystonia. We believe that this study provided a path to unravel candidate genome variants as modifiers. Our findings not only echo the previous research highlighting the defect of mechanosensing and mechanotransduction regulated by TOR1A [48], but provide knowledge for further understanding the disease origin of the DYT1 dystonia as well. We will recommend the physicians to test these variants once the TOR1A ΔE mutation patient show normal alleles within other *TOR1A* locus and other major binding proteins in their study. We also provide a list of candidate genes and genome variants for future mechanistic studies on DYT1 dystonia.

## Supporting information

Table S1 and Figure S1.

## Acknowledgments

We would like to thank the first patient and his family members who provided the DNAs, RNAs, and clinical information necessary for this research study. We would also like to thank Dr. Chin-Hsien Lin (Department of Neurology, National Taiwan University Hospital, Taipei, Taiwan) who kindly provided the DNA sample and clinical information of the second patient. This research was funded by Tri-Service General Hospital, grant number TSGH-C108-021 (C.F.H.), TSGH-C108-022 (C.F.H.) and National Institutes of Health GM129374 (G.W.G.L.), Z99-ES999999 (S.P.W.).

## Author contributions

CFH: Conceptualization, Writing-original draft, Resources. GWGL: Visualization, Writing-review & editing. FCL: Methodology, Investigation. CSH: Data curation, Visualization. SMH: Supervision, Writing - review &editing. JSH: Data curation, Supervision. SPW: Supervision, Writing-review & editing, Resources.

