## Supplementary material for "Whole exome sequencing identifies novel DYT1 dystonia-associated genome variants as potential disease modifiers": Table S1

| **Table S1. Functional annotation clustering-1 (264 variants/195genes)** | | | | | |
| --- | --- | --- | --- | --- | --- |
| **Annotation Cluster 1** | **Enrichment Score: 2.2311** | | | | |
| **Category** | **Term** | **Count** | **%** | ***p* value** | **Genes** |
| INTERPRO | IPR013032:  EGF-like, conserved site | 10 | 5.13 | 1.38E-04 | ATRNL1, TNC, ZAN, TNN, C1R, NRG1, NRG2, MUC17, LRP4, VWA2 |
| INTERPRO | IPR000742:  Epidermal growth factor-like domain | 10 | 5.13 | 4.20E-04 | ATRNL1, TNC, ZAN, TNN, C1R, NRG1, NRG2, MUC17, LRP4, VWA2 |
| UP_KEYWORDS | EGF-like domain | 9 | 4.62 | 0.00192857 | ATRNL1, TNC, ZAN, TNN, C1R, NRG1, NRG2, LRP4, VWA2 |
| SMART | SM00181:EGF | 8 | 4.10 | 0.00312238 | ATRNL1, TNC, ZAN, TNN, C1R, NRG1, LRP4, VWA2 |
| UP_SEQ_FEATURE | domain:EGF-like | 4 | 2.05 | 0.04092413 | ZAN, NRG1, NRG2, MUC17 |
| UP_SEQ_FEATURE | domain:EGF-like 2 | 4 | 2.05 | 0.0532148 | ATRNL1, TNC, TNN, VWA2 |
| UP_SEQ_FEATURE | domain:EGF-like 1 | 3 | 1.54 | 0.31685439 | ATRNL1, TNN, VWA2 |
| **Annotation Cluster 2** | **Enrichment Score: 2.1566** | | | | |
| **Category** | **Term** | **Count** | **%** | ***p* value** | **Genes** |
| GOTERM_MF_DIRECT | GO:0000907~sulfonate dioxygenase activity | 3 | 1.54 | 5.16E-04 | HIF1AN, TET2, TET1 |
| GOTERM_MF_DIRECT | GO:0019798~procollagen-proline dioxygenase activity | 3 | 1.54 | 5.16E-04 | HIF1AN, TET2, TET1 |
| GOTERM_MF_DIRECT | GO:0018602~2,4-dichlorophenoxyacetate alpha-ketoglutarate dioxygenase activity | 3 | 1.54 | 5.16E-04 | HIF1AN, TET2, TET1 |
| GOTERM_MF_DIRECT | GO:0034792~hypophosphite dioxygenase activity | 3 | 1.54 | 5.16E-04 | HIF1AN, TET2, TET1 |
| GOTERM_MF_DIRECT | GO:0052635~C-20 gibberellin 2-beta-dioxygenase activity | 3 | 1.54 | 5.16E-04 | HIF1AN, TET2, TET1 |
| GOTERM_MF_DIRECT | GO:0052634~C-19 gibberellin 2-beta-dioxygenase activity | 3 | 1.54 | 5.16E-04 | HIF1AN, TET2, TET1 |
| UP_SEQ_FEATURE | metal ion-binding site:Iron; catalytic | 4 | 2.05 | 0.00167406 | ALOXE3, HIF1AN, TET2, TET1 |
| GOTERM_MF_DIRECT | GO:0043734~DNA-N1-methyladenine dioxygenase activity | 3 | 1.54 | 0.00177257 | HIF1AN, TET2, TET1 |
| GOTERM_BP_DIRECT | GO:0019511~peptidyl-proline hydroxylation | 3 | 1.54 | 0.00486291 | HIF1AN, TET2, TET1 |
| UP_SEQ_FEATURE | binding site:2-oxoglutarate | 3 | 1.54 | 0.00760956 | HIF1AN, TET2, TET1 |
| UP_KEYWORDS | Dioxygenase | 4 | 2.05 | 0.04376936 | ALOXE3, HIF1AN, TET2, TET1 |
| GOTERM_MF_DIRECT | GO:0005506~iron ion binding | 5 | 2.56 | 0.05563094 | CYP1B1, ALOXE3, HIF1AN, CYP2A7, TET1 |
| UP_KEYWORDS | Iron | 6 | 3.08 | 0.19206252 | CYP1B1, ALOXE3, HIF1AN, CYP2A7, TET2, TET1 |
| UP_KEYWORDS | Monooxygenase | 3 | 1.54 | 0.20141295 | CYP1B1, MICAL3, CYP2A7 |
| UP_KEYWORDS | Oxidoreductase | 8 | 4.10 | 0.31018384 | CYP1B1, ALOXE3, HIF1AN, MICAL3, DHRS4L2, CYP2A7, TET2, TET1 |
| GOTERM_BP_DIRECT | GO:0055114~oxidation-reduction process | 7 | 3.59 | 0.51562726 | CYP1B1, ALOXE3, CFAP61, HIF1AN, MICAL3, DHRS4L2, CYP2A7 |
| **Annotation Cluster 3** | **Enrichment Score: 1.3668** | | | | |
| **Category** | **Term** | **Count** | **%** | ***p* value** | **Genes** |
| UP_SEQ_FEATURE | repeat:4 | 8 | 4.10 | 0.00247366 | VEGFC, KRTAP5-7, HRNR, TUBGCP6, KRTAP13-4, PPP1R15A, MUC17, NUP58 |
| UP_SEQ_FEATURE | repeat:1 | 9 | 4.62 | 0.00329909 | VEGFC, KRTAP5-7, HRNR, TUBGCP6, KRTAP13-4, WRN, PPP1R15A, MUC17, NUP58 |
| UP_SEQ_FEATURE | repeat:2 | 9 | 4.62 | 0.00354268 | VEGFC, KRTAP5-7, HRNR, TUBGCP6, KRTAP13-4, WRN, PPP1R15A, MUC17, NUP58 |
| UP_SEQ_FEATURE | repeat:3 | 8 | 4.10 | 0.00587645 | VEGFC, KRTAP5-7, HRNR, TUBGCP6, KRTAP13-4, PPP1R15A, MUC17, NUP58 |
| UP_SEQ_FEATURE | repeat:7 | 5 | 2.56 | 0.03845699 | KRTAP5-7, HRNR, TUBGCP6, MUC17, NUP58 |
| UP_SEQ_FEATURE | repeat:6 | 5 | 2.56 | 0.05230712 | KRTAP5-7, HRNR, TUBGCP6, MUC17, NUP58 |
| UP_SEQ_FEATURE | repeat:5 | 5 | 2.56 | 0.06838305 | KRTAP5-7, HRNR, TUBGCP6, MUC17, NUP58 |
| UP_SEQ_FEATURE | repeat:9 | 4 | 2.05 | 0.08238395 | HRNR, TUBGCP6, MUC17, NUP58 |
| UP_SEQ_FEATURE | repeat:8 | 4 | 2.05 | 0.11491412 | HRNR, TUBGCP6, MUC17, NUP58 |
| UP_SEQ_FEATURE | repeat:14 | 3 | 1.54 | 0.1768888 | HRNR, MUC17, NUP58 |
| UP_SEQ_FEATURE | repeat:13 | 3 | 1.54 | 0.19058498 | HRNR, MUC17, NUP58 |
| UP_SEQ_FEATURE | repeat:12 | 3 | 1.54 | 0.19748605 | HRNR, MUC17, NUP58 |
| UP_SEQ_FEATURE | repeat:11 | 3 | 1.54 | 0.20789335 | HRNR, MUC17, NUP58 |
| UP_SEQ_FEATURE | repeat:10 | 3 | 1.54 | 0.23940709 | HRNR, MUC17, NUP58 |
| **Annotation Cluster 4** | **Enrichment Score: 1.1214** | | | | |
| **Category** | **Term** | **Count** | **%** | ***p* value** | **Genes** |
| UP_SEQ_FEATURE | domain:LIM zinc-binding | 3 | 1.54 | 0.01386048 | NRAP, LIMCH1, MICAL3 |
| GOTERM_MF_DIRECT | GO:0003779~actin binding | 7 | 3.59 | 0.04689784 | SYNE2, NRAP, MYBPC1, LIMCH1, CAPG, MICAL3, DBN1 |
| UP_KEYWORDS | LIM domain | 3 | 1.54 | 0.14770085 | NRAP, LIMCH1, MICAL3 |
| INTERPRO | IPR001781:Zinc finger, LIM-type | 3 | 1.54 | 0.15314016 | NRAP, LIMCH1, MICAL3 |
| SMART | SM00132:LIM | 3 | 1.54 | 0.16807148 | NRAP, LIMCH1, MICAL3 |
| **Annotation Cluster 5** | **Enrichment Score: 1.0970** | | | | |
| **Category** | **Term** | **Count** | **%** | ***p* value** | **Genes** |
| UP_KEYWORDS | Glycoprotein | 63 | 32.31 | 0.0009329 | PVR, LRRC8E, OR1A1, OR1J2, ZAN, PCDHGA8, UNC93A, GPLD1, GP9, APOB, TSPAN10, SERPINE1, RAET1L, NRG1, PIEZO2, NRG2, VWA2, CEACAM20, ATRNL1, CHST3, VEGFC, SIGLEC8, OR4N2, CHSY1, FCRLB, BTNL10, CHGB, FUT6, OR2A25, TNC, FPR1, DHRS4L2, PCDHGC5, C1R, DISP2, CPZ, OR10G3, OR2T33, TNN, GAL3ST1, NUP58, C10ORF25, KLK2, KLB, FLT4, ASIC4, RHBDF1, TET2, OR4K1, TET1, P2RY13, P2RX7, PROM2, OR3A3, SLC7A2, GRIA1, FREM1, TPSAB1, CHRD, LRP4, MUC17, OR56A5, MUC16 |
| UP_SEQ_FEATURE | glycosylation site:N-linked (GlcNAc...) | 58 | 29.74 | 0.00269477 | PVR, LRRC8E, OR1A1, OR1J2, ZAN, PCDHGA8, UNC93A, GPLD1, GP9, APOB, TSPAN10, SERPINE1, RAET1L, NRG1, PIEZO2, NRG2, VWA2, CEACAM20, ATRNL1, CHST3, VEGFC, SIGLEC8, OR4N2, CHSY1, FCRLB, OR2A25, TNC, FUT6, FPR1, DHRS4L2, PCDHGC5, C1R, DISP2, CPZ, OR10G3, OR2T33, TNN, GAL3ST1, C10ORF25, KLB, KLK2, FLT4, ASIC4, RHBDF1, OR4K1, P2RY13, P2RX7, PROM2, OR3A3, SLC7A2, GRIA1, FREM1, TPSAB1, CHRD, LRP4, MUC17, OR56A5, MUC16 |
| UP_SEQ_FEATURE | disulfide bond | 40 | 20.51 | 0.01638543 | PVR, OR1A1, OR1J2, OR2A25, TNC, ZAN, FPR1, DEFB128, C1R, OR10G3, CPZ, APOB, OR2T33, RAET1L, TNN, NRG1, NRG2, VWA2, CEACAM20, ATRNL1, KLK2, ASIC4, FLT4, OR4K1, NME8, VEGFC, P2RY13, OR3A3, SIGLEC8, FREM1, CX3CR1, OR4N2, IRF3, FCRLB, TPSAB1, CHGB, LRP4, MUC17, OR56A5, MUC16 |
| UP_KEYWORDS | Disulfide bond | 44 | 22.56 | 0.02815672 | PVR, OR1A1, OR1J2, TNC, OR2A25, ZAN, FPR1, DEFB128, C1R, GP9, CPZ, OR10G3, APOB, OR2T33, RAET1L, TNN, NRG1, NRG2, VWA2, CEACAM20, ATRNL1, KLK2, ASIC4, FLT4, OR4K1, NME8, VEGFC, P2RY13, P2RX7, OR3A3, GRIA1, SIGLEC8, FREM1, CX3CR1, OR4N2, IRF3, FCRLB, TPSAB1, BTNL10, CHGB, LRP4, MUC17, OR56A5, MUC16 |
| UP_SEQ_FEATURE | topological domain:Cytoplasmic | 42 | 21.54 | 0.07456026 | PVR, OR1A1, OR1J2, FUT6, OR2A25, ZAN, PCDHGA8, FPR1, PCDHGC5, ATP10D, OR10G3, GP9, OR2T33, NRG1, NRG2, GAL3ST1, CEACAM20, CAMLG, KLB, ATRNL1, ASIC4, FLT4, RHBDF1, CHST3, OR4K1, AQP10, P2RY13, P2RX7, PROM2, SYNE2, OR3A3, SLC7A2, GRIA1, SIGLEC8, CX3CR1, OR4N2, CHSY1, GPAM, MUC17, OR56A5, LRP4, MUC16 |
| UP_SEQ_FEATURE | topological domain:Extracellular | 35 | 17.95 | 0.07487105 | PVR, OR1A1, OR1J2, OR2A25, ZAN, PCDHGA8, FPR1, PCDHGC5, ATP10D, OR10G3, GP9, OR2T33, NRG1, NRG2, CEACAM20, CAMLG, KLB, ATRNL1, FLT4, ASIC4, AQP10, OR4K1, P2RY13, P2RX7, PROM2, OR3A3, SLC7A2, GRIA1, SIGLEC8, CX3CR1, OR4N2, MUC17, OR56A5, LRP4, MUC16 |
| UP_KEYWORDS | Cell membrane | 37 | 18.97 | 0.13379528 | PVR, LRRC8E, PTOV1, OR1A1, OR1J2, OR2A25, ZAN, PCDHGA8, FPR1, UNC93A, PCDHGC5, ATP10D, OR10G3, SMAP1, DYNC2H1, OR2T33, RAET1L, NRG1, TRIP10, NRG2, NLRP6, KLB, FLT4, OR4K1, P2RY13, P2RX7, TNS2, PROM2, SYNE2, OR3A3, SLC7A2, GRIA1, CX3CR1, OR4N2, MUC17, OR56A5, MUC16 |
| UP_KEYWORDS | Transmembrane helix | 60 | 30.77 | 0.19067843 | PVR, LRRC8E, OR1A1, OR1J2, ZAN, PCDHGA8, UNC93A, ATP10D, SPATA31C1, ANKLE1, TSNARE1, GP9, TSPAN10, NRG1, PIEZO2, NRG2, CEACAM20, ATRNL1, CHST3, MYCN, SIGLEC8, CX3CR1, TEX38, OR4N2, CYP2A7, CHSY1, BTNL10, GPAM, CYP1B1, SLC35E2B, FUT6, OR2A25, FPR1, PCDHGC5, DISP2, OR10G3, MTCH2, OR2T33, GAL3ST1, AATK, CAMLG, KLB, FLT4, ASIC4, RHBDF1, OR4K1, AQP10, P2RY13, P2RX7, SYNE2, PROM2, OR3A3, SLC7A2, GRIA1, SLC25A10, RHBDL3, LRP4, MUC17, OR56A5, MUC16 |
| UP_KEYWORDS | Transmembrane | 60 | 30.77 | 0.1978893 | PVR, LRRC8E, OR1A1, OR1J2, ZAN, PCDHGA8, UNC93A, ATP10D, SPATA31C1, ANKLE1, TSNARE1, GP9, TSPAN10, NRG1, PIEZO2, NRG2, CEACAM20, ATRNL1, CHST3, MYCN, SIGLEC8, CX3CR1, TEX38, OR4N2, CYP2A7, CHSY1, BTNL10, GPAM, CYP1B1, SLC35E2B, FUT6, OR2A25, FPR1, PCDHGC5, DISP2, OR10G3, MTCH2, OR2T33, GAL3ST1, AATK, CAMLG, KLB, FLT4, ASIC4, RHBDF1, OR4K1, AQP10, P2RY13, P2RX7, SYNE2, PROM2, OR3A3, SLC7A2, GRIA1, SLC25A10, RHBDL3, LRP4, MUC17, OR56A5, MUC16 |
| UP_SEQ_FEATURE | transmembrane region | 54 | 27.69 | 0.23040636 | PVR, LRRC8E, OR1A1, OR1J2, ZAN, PCDHGA8, UNC93A, ATP10D, TSNARE1, GP9, TSPAN10, NRG1, NRG2, PIEZO2, CEACAM20, ATRNL1, CHST3, SIGLEC8, CX3CR1, TEX38, OR4N2, CHSY1, GPAM, SLC35E2B, OR2A25, FUT6, FPR1, PCDHGC5, DISP2, OR10G3, MTCH2, OR2T33, GAL3ST1, AATK, CAMLG, KLB, FLT4, ASIC4, RHBDF1, OR4K1, AQP10, P2RY13, P2RX7, SYNE2, PROM2, OR3A3, SLC7A2, GRIA1, SLC25A10, RHBDL3, LRP4, MUC17, OR56A5, MUC16 |
| GOTERM_CC_DIRECT | GO:0016021~integral component of membrane | 56 | 28.72 | 0.24611411 | PVR, LRRC8E, OR1A1, OR1J2, ZAN, PCDHGA8, UNC93A, ATP10D, SPATA31C1, ANKLE1, TSNARE1, TSPAN10, NRG1, NRG2, PIEZO2, CEACAM20, ATRNL1, CHST3, MYCN, SIGLEC8, CX3CR1, TEX38, OR4N2, CYP2A7, CHSY1, BTNL10, GPAM, CYP1B1, SLC35E2B, OR2A25, FUT6, FPR1, PCDHGC5, DISP2, OR10G3, MTCH2, OR2T33, INPP5B, GAL3ST1, AATK, CAMLG, KLB, RHBDF1, OR4K1, P2RY13, SYNE2, PROM2, OR3A3, GRIA1, FREM1, SLC25A10, RHBDL3, LRP4, MUC17, OR56A5, MUC16 |
| GOTERM_CC_DIRECT | GO:0005886~plasma membrane | 45 | 23.08 | 0.27578142 | PVR, LRRC8E, PTOV1, OR1A1, OR1J2, OR2A25, ZAN, PCDHGA8, FPR1, UNC93A, PCDHGC5, ATP10D, DISP2, GP9, OR10G3, PLCL1, METTL22, APOB, SMAP1, SERPINE1, OR2T33, DYNC2H1, AHNAK2, RAET1L, INPP5B, NRG2, NLRP6, KLB, FLT4, MICAL3, OR4K1, CAPN3, AQP10, P2RY13, P2RX7, TNS2, OR3A3, GRIA1, SLC7A2, CX3CR1, OR4N2, DBN1, GPAM, OR56A5, MUC16 |
| UP_KEYWORDS | Membrane | 76 | 38.97 | 0.27994864 | PVR, LRRC8E, OR1A1, OR1J2, ZAN, PCDHGA8, UNC93A, ATP10D, SPATA31C1, ANKLE1, TSNARE1, GP9, SMAP1, TSPAN10, DYNC2H1, RAET1L, NRG1, VPS11, PIEZO2, NRG2, CEACAM20, NLRP6, ATRNL1, CHST3, MYCN, TNS2, SIGLEC8, CX3CR1, TEX38, AKAP6, OR4N2, CHSY1, CYP2A7, PPP1R15A, BTNL10, GPAM, PTOV1, CYP1B1, SLC35E2B, FUT6, OR2A25, FPR1, AKAP13, PCDHGC5, DISP2, OR10G3, DOCK1, MTCH2, OR2T33, TRIP10, INPP5B, GAL3ST1, NUP58, AATK, CAMLG, KLB, ASIC4, FLT4, RHBDF1, OR4K1, AQP10, P2RY13, P2RX7, PROM2, SYNE2, OR3A3, GRIA1, SLC7A2, SLC25A10, PLA2G4F, HGS, RHBDL3, LRP4, MUC17, OR56A5, MUC16 |
| UP_KEYWORDS | Signal | 41 | 21.03 | 0.47247256 | PVR, SLC35E2B, TNC, ZAN, PCDHGA8, DEFB128, DHRS4L2, GPLD1, PCDHGC5, C1R, GP9, CPZ, APOB, SERPINE1, RAET1L, TNN, NRG1, PIEZO2, NRG2, VWA2, C10ORF25, CEACAM20, ATRNL1, KLK2, FLT4, VEGFC, PROM2, GRIA1, SIGLEC8, FREM1, PLA2G4F, GHRL, CYP2A7, C7ORF57, FCRLB, TPSAB1, CHGB, BTNL10, CHRD, LRP4, MUC17 |
| UP_SEQ_FEATURE | signal peptide | 33 | 16.92 | 0.51903421 | PVR, TNC, ZAN, PCDHGA8, DEFB128, DHRS4L2, GPLD1, PCDHGC5, C1R, GP9, CPZ, APOB, SERPINE1, RAET1L, TNN, VWA2, C10ORF25, CEACAM20, ATRNL1, KLK2, FLT4, VEGFC, PROM2, GRIA1, SIGLEC8, FREM1, GHRL, FCRLB, TPSAB1, CHGB, CHRD, MUC17, LRP4 |
| **Annotation Cluster 6** | **Enrichment Score: 1.0496** | | | | |
| **Category** | **Term** | **Count** | **%** | ***p* value** | **Genes** |
| KEGG_PATHWAY | hsa04740:Olfactory transduction | 9 | 4.62 | 0.03060606 | OR3A3, OR1A1, OR1J2, OR2A25, OR2T33, OR4N2, OR4K1, OR56A5, OR10G3 |
| GOTERM_MF_DIRECT | GO: 0004984~olfactory receptor activity | 9 | 4.62 | 0.05001473 | OR3A3, OR1A1, OR1J2, OR2A25, OR2T33, OR4N2, OR4K1, OR56A5, OR10G3 |
| GOTERM_BP_DIRECT | GO:0007186~G-protein coupled receptor signaling pathway | 15 | 7.69 | 0.05485615 | NLRP6, OR1A1, ATRNL1, OR3A3, OR1J2, OR2A25, CX3CR1, FPR1, OR2T33, AKAP13, GHRL, OR4N2, OR4K1, OR56A5, OR10G3 |
| GOTERM_BP_DIRECT | GO:0050911~detection of chemical stimulus involved in sensory perception of smell | 9 | 4.62 | 0.05566192 | OR3A3, OR1A1, OR1J2, OR2A25, OR2T33, OR4N2, OR4K1, OR56A5, OR10G3 |
| INTERPRO | IPR000725:Olfactory receptor | 9 | 4.62 | 0.05728121 | OR3A3, OR1A1, OR1J2, OR2A25, OR2T33, OR4N2, OR4K1, OR56A5, OR10G3 |
| UP_KEYWORDS | Olfaction | 9 | 4.62 | 0.05992948 | OR3A3, OR1A1, OR1J2, OR2A25, OR2T33, OR4N2, OR4K1, OR56A5, OR10G3 |
| UP_KEYWORDS | Sensory transduction | 11 | 5.64 | 0.06338104 | OR3A3, OR1A1, OR1J2, OR2A25, HPS1, OR2T33, OR4N2, OR4K1, PIEZO2, OR56A5, OR10G3 |
| INTERPRO | IPR000276:G protein-coupled receptor, rhodopsin-like | 12 | 6.15 | 0.08874814 | P2RY13, OR3A3, OR1A1, OR1J2, OR2A25, CX3CR1, FPR1, OR2T33, OR4N2, OR4K1, OR56A5, OR10G3 |
| INTERPRO | IPR017452:GPCR, rhodopsin-like, 7TM | 12 | 6.15 | 0.10027694 | P2RY13, OR3A3, OR1A1, OR1J2, OR2A25, CX3CR1, FPR1, OR2T33, OR4N2, OR4K1, OR56A5, OR10G3 |
| UP_KEYWORDS | Transducer | 13 | 6.67 | 0.14329892 | PLCL1, P2RY13, OR1A1, OR3A3, OR1J2, OR2A25, CX3CR1, FPR1, OR2T33, OR4N2, OR4K1, OR56A5, OR10G3 |
| UP_KEYWORDS | G-protein coupled receptor | 12 | 6.15 | 0.17266455 | P2RY13, OR3A3, OR1A1, OR1J2, OR2A25, CX3CR1, FPR1, OR2T33, OR4N2, OR4K1, OR56A5, OR10G3 |
| UP_KEYWORDS | Receptor | 19 | 9.74 | 0.29155137 | PVR, OR1A1, OR1J2, OR2A25, FLT4, FPR1, OR4K1, OR10G3, P2RY13, P2RX7, OR3A3, GRIA1, CX3CR1, OR2T33, OR4N2, TRIP10, LRP4, OR56A5, TRAF3 |
| GOTERM_MF_DIRECT | GO:0004930~G-protein coupled receptor activity | 9 | 4.62 | 0.34715936 | OR3A3, OR1A1, OR1J2, OR2A25, OR2T33, OR4N2, OR4K1, OR56A5, OR10G3 |
| **Annotation Cluster 7** | **Enrichment Score: 1.0343** | | | | |
| **Category** | **Term** | **Count** | **%** | ***p* value** | **Genes** |
| GOTERM_MF_DIRECT | GO:0005089~Rho guanyl-nucleotide exchange factor activity | 4 | 2.05 | 0.03557909 | ARHGEF3, DOCK1, ARHGEF37, AKAP13 |
| GOTERM_BP_DIRECT | GO:0043547~positive regulation of GTPase activity | 11 | 5.64 | 0.04904833 | ARHGEF3, DOCK1, SMAP1, KLB, ARHGEF37, HPS1, AKAP13, TRIP10, NRG1, INPP5B, NRG2 |
| UP_KEYWORDS | Guanine-nucleotide releasing factor | 5 | 2.56 | 0.05256756 | ARHGEF3, DOCK1, ARHGEF37, HPS1, AKAP13 |
| GOTERM_MF_DIRECT | GO:0005085~guanyl-nucleotide exchange factor activity | 4 | 2.05 | 0.09898216 | ARHGEF3, DOCK1, HPS1, AKAP13 |
| UP_SEQ_FEATURE | domain:DH | 3 | 1.54 | 0.13694347 | ARHGEF3, ARHGEF37, AKAP13 |
| INTERPRO | IPR000219:Dbl homology (DH) domain | 3 | 1.54 | 0.14978632 | ARHGEF3, ARHGEF37, AKAP13 |
| SMART | SM00325:RhoGEF | 3 | 1.54 | 0.15364418 | ARHGEF3, ARHGEF37, AKAP13 |
| GOTERM_BP_DIRECT | GO:0035023~regulation of Rho protein signal transduction | 3 | 1.54 | 0.185707 | ARHGEF3, ARHGEF37, AKAP13 |
| **Annotation Cluster 8** | **Enrichment Score: 0.9803** | | | | |
| **Category** | **Term** | **Count** | **%** | ***p* value** | **Genes** |
| GOTERM_MF_DIRECT | GO:0003779~actin binding | 7 | 3.59 | 0.04689784 | SYNE2, NRAP, MYBPC1, LIMCH1, CAPG, MICAL3, DBN1 |
| SMART | SM00033:CH | 3 | 1.54 | 0.14652756 | SYNE2, LIMCH1, MICAL3 |
| INTERPRO | IPR001715:Calponin homology domain | 3 | 1.54 | 0.16669639 | SYNE2, LIMCH1, MICAL3 |
| **Annotation Cluster 9** | **Enrichment Score: 0.9678** | | | | |
| **Category** | **Term** | **Count** | **%** | ***p* value** | **Genes** |
| INTERPRO | IPR023795:Protease inhibitor I4, serpin, conserved site | 3 | 1.54 | 0.04017305 | SERPINB6, SERPINE1, SERPINB11 |
| INTERPRO | IPR023796:Serpin domain | 3 | 1.54 | 0.05185145 | SERPINB6, SERPINE1, SERPINB11 |
| INTERPRO | IPR000215:Serpin family | 3 | 1.54 | 0.05185145 | SERPINB6, SERPINE1, SERPINB11 |
| SMART | SM00093:SERPIN | 3 | 1.54 | 0.05485951 | SERPINB6, SERPINE1, SERPINB11 |
| UP_SEQ_FEATURE | site:Reactive bond | 3 | 1.54 | 0.06837429 | SERPINB6, SERPINE1, SERPINB11 |
| UP_KEYWORDS | Serine protease inhibitor | 3 | 1.54 | 0.19456279 | SERPINB6, SERPINE1, SERPINB11 |
| GOTERM_MF_DIRECT | GO:0004867~serine-type endopeptidase inhibitor activity | 3 | 1.54 | 0.2301332 | SERPINB6, SERPINE1, SERPINB11 |
| UP_KEYWORDS | Protease inhibitor | 3 | 1.54 | 0.32654821 | SERPINB6, SERPINE1, SERPINB11 |
| GOTERM_BP_DIRECT | GO:0010951~negative regulation of endopeptidase activity | 3 | 1.54 | 0.32867818 | SERPINB6, SERPINE1, SERPINB11 |
| **Annotation Cluster 10** | **Enrichment Score: 0.9676** | | | | |
| **Category** | **Term** | **Count** | **%** | ***p* value** | **Genes** |
| INTERPRO | IPR003598:Immunoglobulin subtype 2 | 7 | 3.59 | 0.03449055 | CEACAM20, IGFN1, MYBPC1, SIGLEC8, FLT4, NRG1, NRG2 |
| SMART | SM00408:IGc2 | 7 | 3.59 | 0.0431385 | CEACAM20, IGFN1, MYBPC1, SIGLEC8, FLT4, NRG1, NRG2 |
| UP_SEQ_FEATURE | domain:Ig-like C2-type 1 | 6 | 3.08 | 0.04427673 | PVR, CEACAM20, MYBPC1, SIGLEC8, FLT4, FCRLB |
| UP_SEQ_FEATURE | domain:Ig-like C2-type 2 | 6 | 3.08 | 0.04506012 | PVR, CEACAM20, MYBPC1, SIGLEC8, FLT4, FCRLB |
| INTERPRO | IPR013098:Immunoglobulin I-set | 5 | 2.56 | 0.04778435 | IGFN1, MYBPC1, FLT4, NRG1, NRG2 |
| UP_KEYWORDS | Immunoglobulin domain | 10 | 5.13 | 0.07442334 | PVR, CEACAM20, IGFN1, MYBPC1, SIGLEC8, FLT4, FCRLB, NRG1, BTNL10, NRG2 |
| INTERPRO | IPR003599:Immunoglobulin subtype | 9 | 4.62 | 0.10881217 | PVR, CEACAM20, IGFN1, MYBPC1, SIGLEC8, FLT4, FCRLB, NRG1, NRG2 |
| SMART | SM00409:IG | 9 | 4.62 | 0.13656689 | PVR, CEACAM20, IGFN1, MYBPC1, SIGLEC8, FLT4, FCRLB, NRG1, NRG2 |
| UP_SEQ_FEATURE | domain:Ig-like C2-type 4 | 3 | 1.54 | 0.16336033 | CEACAM20, MYBPC1, FLT4 |
| INTERPRO | IPR013783:Immunoglobulin-like fold | 13 | 6.67 | 0.20818552 | PVR, CEACAM20, MYBPC1, TNC, FLT4, IGFN1, GBE1, SIGLEC8, TNN, FCRLB, NRG1, BTNL10, NRG2 |
| INTERPRO | IPR007110:Immunoglobulin-like domain | 10 | 5.13 | 0.32721003 | PVR, CEACAM20, IGFN1, MYBPC1, SIGLEC8, FLT4, FCRLB, NRG1, BTNL10, NRG2 |
| UP_SEQ_FEATURE | domain:Ig-like C2-type 3 | 3 | 1.54 | 0.3446877 | CEACAM20, MYBPC1, FLT4 |
| GOTERM_BP_DIRECT | GO:0018108~peptidyl-tyrosine phosphorylation | 3 | 1.54 | 0.43870861 | FLT4, NRG1, NRG2 |
| **Annotation Cluster 11** | **Enrichment Score: 0.9212** | | | | |
| **Category** | **Term** | **Count** | **%** | ***p* value** | **Genes** |
| SMART | SM00181:EGF | 8 | 4.10 | 0.00312238 | ATRNL1, TNC, ZAN, TNN, C1R, NRG1, LRP4, VWA2 |
| INTERPRO | IPR001881:EGF-like calcium-binding | 3 | 1.54 | 0.34701215 | C1R, LRP4, VWA2 |
| SMART | SM00179:EGF_CA | 3 | 1.54 | 0.37462283 | C1R, LRP4, VWA2 |
| GOTERM_MF_DIRECT | GO:0005509~calcium ion binding | 8 | 4.10 | 0.50926212 | HRNR, PCDHGA8, PCDHGC5, C1R, RHBDL3, CAPN3, LRP4, VWA2 |
| **Annotation Cluster 12** | **Enrichment Score: 0.8643** | | | | |
| **Category** | **Term** | **Count** | **%** | ***p* value** | **Genes** |
| UP_SEQ_FEATURE | domain:Fibronectin type-III 3 | 4 | 2.05 | 0.04618849 | IGFN1, MYBPC1, TNC, TNN |
| UP_SEQ_FEATURE | domain:Fibronectin type-III 4 | 3 | 1.54 | 0.12097393 | IGFN1, TNC, TNN |
| UP_SEQ_FEATURE | domain:Fibronectin type-III 2 | 4 | 2.05 | 0.13380932 | IGFN1, MYBPC1, TNC, TNN |
| UP_SEQ_FEATURE | domain:Fibronectin type-III 1 | 4 | 2.05 | 0.13597236 | IGFN1, MYBPC1, TNC, TNN |
| SMART | SM00060:FN3 | 4 | 2.05 | 0.19877763 | IGFN1, MYBPC1, TNC, TNN |
| INTERPRO | IPR003961:Fibronectin, type III | 4 | 2.05 | 0.32260263 | IGFN1, MYBPC1, TNC, TNN |
| **Annotation Cluster 13** | **Enrichment Score: 0.7570** | | | | |
| **Category** | **Term** | **Count** | **%** | ***p* value** | **Genes** |
| GOTERM_BP_DIRECT | GO:0035666~TRIF-dependent toll-like receptor signaling pathway | 3 | 1.54 | 0.03001222 | IRF3, CHUK, TRAF3 |
| KEGG_PATHWAY | hsa04622:RIG-I-like receptor signaling pathway | 3 | 1.54 | 0.13786424 | IRF3, CHUK, TRAF3 |
| KEGG_PATHWAY | hsa05168:Herpes simplex infection | 4 | 2.05 | 0.24196398 | TAF5, IRF3, CHUK, TRAF3 |
| KEGG_PATHWAY | hsa04620:Toll-like receptor signaling pathway | 3 | 1.54 | 0.25906033 | IRF3, CHUK, TRAF3 |
| KEGG_PATHWAY | hsa05169:Epstein-Barr virus infection | 3 | 1.54 | 0.3144158 | IRF3, CHUK, TRAF3 |
| KEGG_PATHWAY | hsa05160:Hepatitis C | 3 | 1.54 | 0.35203734 | IRF3, CHUK, TRAF3 |
| **Annotation Cluster 14** | **Enrichment Score: 0.6739** | | | | |
| **Category** | **Term** | **Count** | **%** | ***p* value** | **Genes** |
| UP_SEQ_FEATURE | repeat:LRR 9 | 4 | 2.05 | 0.13597236 | LRRC8E, SYNE2, LRRC2, MUC16 |
| UP_SEQ_FEATURE | repeat:LRR 13 | 3 | 1.54 | 0.16000831 | LRRC8E, SYNE2, MUC16 |
| UP_SEQ_FEATURE | repeat:LRR 3 | 6 | 3.08 | 0.16184808 | LRRC8E, NLRP6, SYNE2, LRRC2, PRAMEF1, MUC16 |
| UP_SEQ_FEATURE | repeat:LRR 8 | 4 | 2.05 | 0.16831037 | LRRC8E, SYNE2, LRRC2, MUC16 |
| UP_SEQ_FEATURE | repeat:LRR 5 | 5 | 2.56 | 0.17862185 | LRRC8E, NLRP6, SYNE2, LRRC2, MUC16 |
| UP_SEQ_FEATURE | repeat:LRR 1 | 6 | 3.08 | 0.19208783 | LRRC8E, NLRP6, SYNE2, LRRC2, PRAMEF1, MUC16 |
| UP_SEQ_FEATURE | repeat:LRR 2 | 6 | 3.08 | 0.1937351 | LRRC8E, NLRP6, SYNE2, LRRC2, PRAMEF1, MUC16 |
| UP_SEQ_FEATURE | repeat:LRR 12 | 3 | 1.54 | 0.21137551 | LRRC8E, SYNE2, MUC16 |
| UP_SEQ_FEATURE | repeat:LRR 4 | 5 | 2.56 | 0.22582289 | LRRC8E, NLRP6, SYNE2, LRRC2, MUC16 |
| UP_SEQ_FEATURE | repeat:LRR 7 | 4 | 2.05 | 0.23665721 | LRRC8E, SYNE2, LRRC2, MUC16 |
| UP_SEQ_FEATURE | repeat:LRR 11 | 3 | 1.54 | 0.24997281 | LRRC8E, SYNE2, MUC16 |
| UP_SEQ_FEATURE | repeat:LRR 10 | 3 | 1.54 | 0.30283358 | LRRC8E, SYNE2, MUC16 |
| UP_SEQ_FEATURE | repeat:LRR 6 | 4 | 2.05 | 0.32361439 | LRRC8E, SYNE2, LRRC2, MUC16 |
| UP_KEYWORDS | Leucine-rich repeat | 5 | 2.56 | 0.33690863 | LRRC8E, NLRP6, LRRC2, PRAMEF1, GP9 |
| **Annotation Cluster 15** | **Enrichment Score: 0.6259** | | | | |
| **Category** | **Term** | **Count** | **%** | ***p* value** | **Genes** |
| GOTERM_MF_DIRECT | GO:0046934~phosphatidylinositol-4,5-bisphosphate 3-kinase activity | 3 | 1.54 | 0.11455346 | KLB, NRG1, NRG2 |
| GOTERM_BP_DIRECT | GO:0014066~regulation of phosphatidylinositol 3-kinase signaling | 3 | 1.54 | 0.1752574 | KLB, NRG1, NRG2 |
| GOTERM_BP_DIRECT | GO:0046854~phosphatidylinositol phosphorylation | 3 | 1.54 | 0.23181768 | KLB, NRG1, NRG2 |
| GOTERM_BP_DIRECT | GO:0048015~phosphatidylinositol-mediated signaling | 3 | 1.54 | 0.27495752 | KLB, NRG1, NRG2 |
| GOTERM_MF_DIRECT | GO:0005088~Ras guanyl-nucleotide exchange factor activity | 3 | 1.54 | 0.29251815 | KLB, NRG1, NRG2 |
| GOTERM_BP_DIRECT | GO:0000165~MAPK cascade | 4 | 2.05 | 0.46908184 | CUL3, KLB, NRG1, NRG2 |
| **Annotation Cluster 16** | **Enrichment Score: 0.5092** | | | | |
| **Category** | **Term** | **Count** | **%** | ***p* value** | **Genes** |
| UP_KEYWORDS | Serine protease | 4 | 2.05 | 0.14834205 | KLK2, C1R, RHBDL3, TPSAB1 |
| GOTERM_MF_DIRECT | GO:0004252~serine-type endopeptidase activity | 5 | 2.56 | 0.21690111 | KLK2, RHBDF1, C1R, RHBDL3, TPSAB1 |
| UP_KEYWORDS | Protease | 8 | 4.10 | 0.2541185 | KLK2, RHBDF1, USP36, C1R, RHBDL3, TPSAB1, CAPN3, CPZ |
| UP_SEQ_FEATURE | domain:Peptidase S1 | 3 | 1.54 | 0.2781913 | KLK2, C1R, TPSAB1 |
| INTERPRO | IPR001314:Peptidase S1A, chymotrypsin-type | 3 | 1.54 | 0.29750862 | KLK2, C1R, TPSAB1 |
| INTERPRO | IPR001254:Peptidase S1 | 3 | 1.54 | 0.32237061 | KLK2, C1R, TPSAB1 |
| SMART | SM00020:Tryp_SPc | 3 | 1.54 | 0.34125495 | KLK2, C1R, TPSAB1 |
| INTERPRO | IPR009003:Trypsin-like cysteine/serine peptidase domain | 3 | 1.54 | 0.3505097 | KLK2, C1R, TPSAB1 |
| GOTERM_BP_DIRECT | GO:0006508~proteolysis | 6 | 3.08 | 0.53650166 | KLK2, RHBDF1, C1R, TPSAB1, CAPN3, CPZ |
| UP_SEQ_FEATURE | active site:Charge relay system | 3 | 1.54 | 0.57791771 | KLK2, C1R, TPSAB1 |
| **Annotation Cluster 17** | **Enrichment Score: 0.3366** | | | | |
| **Category** | **Term** | **Count** | **%** | ***p* value** | **Genes** |
| GOTERM_CC_DIRECT | GO:0005913~cell-cell adherens junction | 5 | 2.56 | 0.38856203 | PVR, CAPG, GLOD4, UBAP2, DBN1 |
| GOTERM_BP_DIRECT | GO:0098609~cell-cell adhesion | 4 | 2.05 | 0.49126598 | CAPG, GLOD4, UBAP2, DBN1 |
| GOTERM_MF_DIRECT | GO:0098641~cadherin binding involved in cell-cell adhesion | 4 | 2.05 | 0.51218869 | CAPG, GLOD4, UBAP2, DBN1 |
| **Annotation Cluster 18** | **Enrichment Score: 0.2833** | | | | |
| **Category** | **Term** | **Count** | **%** | ***p* value** | **Genes** |
| UP_KEYWORDS | Mitochondrion | 13 | 6.67 | 0.36509064 | CYP1B1, MRPL12, SYNE2, MTCH2, SLC25A10, TRMT10C, GLOD4, LIPT2, RBFA, GPAM, PPP1R15A, TRAF3, MRM3 |
| UP_KEYWORDS | Transit peptide | 6 | 3.08 | 0.57192648 | MRPL12, TRMT10C, LIPT2, RBFA, GPAM, MRM3 |
| UP_SEQ_FEATURE | transit peptide:Mitochondrion | 5 | 2.56 | 0.67683911 | MRPL12, TRMT10C, LIPT2, RBFA, GPAM |
| **Annotation Cluster 19** | **Enrichment Score: 0.2698** | | | | |
| **Category** | **Term** | **Count** | **%** | ***p* value** | **Genes** |
| GOTERM_BP_DIRECT | GO:0035556~intracellular signal transduction | 7 | 3.59 | 0.19838688 | PLCL1, ARHGEF3, TNS2, AKAP6, AKAP13, NRG1, NRG2 |
| INTERPRO | IPR011993:Pleckstrin homology-like domain | 5 | 2.56 | 0.59279396 | PLCL1, ARHGEF3, TNS2, AKAP13, MTMR6 |
| UP_SEQ_FEATURE | domain:PH | 3 | 1.54 | 0.67983768 | PLCL1, ARHGEF3, AKAP13 |
| INTERPRO | IPR001849:Pleckstrin homology domain | 3 | 1.54 | 0.73902241 | PLCL1, ARHGEF3, AKAP13 |
| SMART | SM00233:PH | 3 | 1.54 | 0.75768944 | PLCL1, ARHGEF3, AKAP13 |
| **Annotation Cluster 20** | **Enrichment Score: 0.2653** | | | | |
| **Category** | **Term** | **Count** | **%** | ***p* value** | **Genes** |
| INTERPRO | IPR011992:EF-hand-like domain | 4 | 2.05 | 0.50628352 | PLCL1, HRNR, RHBDL3, CAPN3 |
| GOTERM_MF_DIRECT | GO:0005509~calcium ion binding | 8 | 4.10 | 0.50926212 | HRNR, PCDHGA8, PCDHGC5, C1R, RHBDL3, CAPN3, LRP4, VWA2 |
| UP_SEQ_FEATURE | domain:EF-hand 2 | 3 | 1.54 | 0.53095765 | HRNR, RHBDL3, CAPN3 |
| UP_SEQ_FEATURE | domain:EF-hand 1 | 3 | 1.54 | 0.53095765 | HRNR, RHBDL3, CAPN3 |
| INTERPRO | IPR002048:EF-hand domain | 3 | 1.54 | 0.64841304 | HRNR, RHBDL3, CAPN3 |
| **Annotation Cluster 21** | **Enrichment Score: 0.2340** | | | | |
| **Category** | **Term** | **Count** | **%** | ***p* value** | **Genes** |
| GOTERM_CC_DIRECT | GO:0005768~endosome | 4 | 2.05 | 0.37211851 | HGS, DOPEY2, VPS11, TRAF3 |
| INTERPRO | IPR013083:Zinc finger, RING/FYVE/PHD-type | 5 | 2.56 | 0.63928797 | JADE2, HGS, VPS11, RNF213, TRAF3 |
| UP_KEYWORDS | Endosome | 4 | 2.05 | 0.83481364 | HGS, VPS11, INPP5B, TRAF3 |
| **Annotation Cluster 22** | **Enrichment Score: 0.2338** | | | | |
| **Category** | **Term** | **Count** | **%** | ***p* value** | **Genes** |
| GOTERM_CC_DIRECT | GO:0000139~Golgi membrane | 8 | 4.10 | 0.35637986 | CUL3, GRIA1, FUT6, RHBDF1, CHST3, DOPEY2, CHSY1, GAL3ST1 |
| UP_SEQ_FEATURE | topological domain:Lumenal | 5 | 2.56 | 0.62369864 | FUT6, RHBDF1, CHST3, CHSY1, GAL3ST1 |
| UP_KEYWORDS | Golgi apparatus | 8 | 4.10 | 0.64802668 | CUL3, FUT6, RHBDF1, CHST3, CHSY1, TRIP10, INPP5B, GAL3ST1 |
| UP_KEYWORDS | Signal-anchor | 4 | 2.05 | 0.8058259 | FUT6, CHST3, CHSY1, GAL3ST1 |
| **Annotation Cluster 23** | **Enrichment Score: 0.2247** | | | | |
| **Category** | **Term** | **Count** | **%** | ***p* value** | **Genes** |
| UP_SEQ_FEATURE | repeat:ANK 1 | 4 | 2.05 | 0.41091593 | POTEF, ANKLE1, POTEE, MUC16 |
| UP_SEQ_FEATURE | repeat:ANK 2 | 4 | 2.05 | 0.41284028 | POTEF, ANKLE1, POTEE, MUC16 |
| UP_SEQ_FEATURE | repeat:ANK 3 | 3 | 1.54 | 0.58856794 | POTEF, ANKLE1, POTEE |
| INTERPRO | IPR002110:Ankyrin repeat | 3 | 1.54 | 0.70788032 | POTEF, ANKLE1, POTEE |
| UP_KEYWORDS | ANK repeat | 3 | 1.54 | 0.71376952 | POTEF, ANKLE1, POTEE |
| INTERPRO | IPR020683:Ankyrin repeat-containing domain | 3 | 1.54 | 0.72768842 | POTEF, ANKLE1, POTEE |
| SMART | SM00248:ANK | 3 | 1.54 | 0.7282786 | POTEF, ANKLE1, POTEE |
| **Annotation Cluster 24** | **Enrichment Score: 0.2153** | | | | |
| **Category** | **Term** | **Count** | **%** | ***p* value** | **Genes** |
| UP_SEQ_FEATURE | zinc finger region:C2H2-type 2 | 9 | 4.62 | 0.23310117 | ZSCAN30, ZNF223, ZIC5, KLF17, ZNF414, ZNF221, ZFP28, KLF1, ZNF470 |
| UP_SEQ_FEATURE | zinc finger region:C2H2-type 3 | 9 | 4.62 | 0.26078577 | ZSCAN30, ZNF223, ZIC5, KLF17, ZNF414, ZNF221, ZFP28, KLF1, ZNF470 |
| UP_SEQ_FEATURE | zinc finger region:C2H2-type 1 | 8 | 4.10 | 0.27675301 | ZSCAN30, ZNF223, KLF17, ZNF414, ZNF221, ZFP28, KLF1, ZNF470 |
| UP_KEYWORDS | Zinc | 26 | 13.33 | 0.28862695 | AKAP13, RNF213, CPZ, JADE2, SMAP1, ZNF223, VPS11, TNIP2, ZNF470, TRAF3, MICAL3, KLF17, WRN, ZNF221, TET2, AMPD2, ZFP28, TET1, TNS2, NRAP, ZSCAN30, LIMCH1, ZIC5, HGS, ZNF414, KLF1 |
| UP_KEYWORDS | Metal-binding | 38 | 19.49 | 0.33352475 | HRNR, CYP1B1, ALOXE3, AKAP13, ATP10D, C1R, RNF213, CPZ, JADE2, SMAP1, HIF1AN, ZNF223, VPS11, TNIP2, INPP5B, ZNF470, TRAF3, MICAL3, KLF17, ZNF221, WRN, ZFP28, TET2, AMPD2, CAPN3, TET1, TNS2, NRAP, ZSCAN30, LIMCH1, FREM1, ZIC5, PLA2G4F, HGS, CHSY1, CYP2A7, ZNF414, KLF1 |
| GOTERM_MF_DIRECT | GO:0003700~transcription factor activity, sequence-specific DNA binding | 11 | 5.64 | 0.4117819 | TAF5, ZSCAN30, ZNF223, KLF17, IRF3, ZNF221, ZFP28, KLF1, ZNF470, TCF3, MYCN |
| UP_SEQ_FEATURE | zinc finger region:C2H2-type 14 | 3 | 1.54 | 0.44734353 | ZNF221, ZFP28, ZNF470 |
| INTERPRO | IPR015880:Zinc finger, C2H2-like | 9 | 4.62 | 0.45450212 | ZSCAN30, ZNF223, ZIC5, KLF17, ZNF414, ZNF221, ZFP28, KLF1, ZNF470 |
| INTERPRO | IPR007087:Zinc finger, C2H2 | 9 | 4.62 | 0.50806115 | ZSCAN30, ZNF223, ZIC5, KLF17, ZNF414, ZNF221, ZFP28, KLF1, ZNF470 |
| UP_KEYWORDS | Zinc-finger | 18 | 9.23 | 0.51555435 | AKAP13, KLF17, ZNF221, ZFP28, RNF213, TET1, JADE2, TNS2, SMAP1, ZSCAN30, ZIC5, HGS, ZNF414, TNIP2, VPS11, ZNF470, KLF1, TRAF3 |
| SMART | SM00355:ZnF_C2H2 | 9 | 4.62 | 0.52447143 | ZSCAN30, ZNF223, ZIC5, KLF17, ZNF414, ZNF221, ZFP28, KLF1, ZNF470 |
| INTERPRO | IPR013087:Zinc finger C2H2-type/integrase DNA-binding domain | 8 | 4.10 | 0.5335161 | ZSCAN30, ZNF223, ZIC5, KLF17, ZNF221, ZFP28, KLF1, ZNF470 |
| GOTERM_MF_DIRECT | GO:0046872~metal ion binding | 20 | 10.26 | 0.57205133 | ALOXE3, AKAP13, KLF17, ZNF221, AMPD2, ZFP28, TNS2, SMAP1, ZSCAN30, FREM1, ZNF223, ZIC5, HGS, PLA2G4F, CHSY1, ZNF414, TNIP2, INPP5B, ZNF470, KLF1 |
| UP_SEQ_FEATURE | zinc finger region:C2H2-type 13 | 3 | 1.54 | 0.58683537 | ZNF221, ZFP28, ZNF470 |
| UP_SEQ_FEATURE | zinc finger region:C2H2-type 7 | 5 | 2.56 | 0.64592722 | ZSCAN30, ZNF223, ZNF221, ZFP28, ZNF470 |
| UP_SEQ_FEATURE | zinc finger region:C2H2-type 12 | 3 | 1.54 | 0.70297602 | ZNF221, ZFP28, ZNF470 |
| UP_SEQ_FEATURE | zinc finger region:C2H2-type 9 | 4 | 2.05 | 0.71276579 | ZNF223, ZNF221, ZFP28, ZNF470 |
| INTERPRO | IPR001909:Krueppel-associated box | 4 | 2.05 | 0.74795666 | ZNF223, ZNF221, ZFP28, ZNF470 |
| SMART | SM00349:KRAB | 4 | 2.05 | 0.77134214 | ZNF223, ZNF221, ZFP28, ZNF470 |
| UP_SEQ_FEATURE | zinc finger region:C2H2-type 5 | 5 | 2.56 | 0.77185574 | ZSCAN30, ZNF223, ZNF221, ZFP28, ZNF470 |
| UP_SEQ_FEATURE | zinc finger region:C2H2-type 8 | 4 | 2.05 | 0.77580815 | ZNF223, ZNF221, ZFP28, ZNF470 |
| UP_SEQ_FEATURE | zinc finger region:C2H2-type 11 | 3 | 1.54 | 0.78358919 | ZNF221, ZFP28, ZNF470 |
| GOTERM_BP_DIRECT | GO:0006355~regulation of transcription, DNA-templated | 13 | 6.67 | 0.80090218 | PTOV1, TAF5, ZNF221, ZFP28, ZSCAN30, CDK11A, ZNF223, CDK11B, IRF3, ZNF414, KLF1, TCF3, ZNF470 |
| UP_SEQ_FEATURE | zinc finger region:C2H2-type 4 | 5 | 2.56 | 0.81464049 | ZSCAN30, ZNF223, ZIC5, ZFP28, ZNF470 |
| UP_KEYWORDS | DNA-binding | 17 | 8.72 | 0.82045289 | KLF17, WRN, ZNF221, ZFP28, TET2, TET1, MYCN, HOXD9, HJURP, ZSCAN30, ZNF223, ZIC5, IRF3, ZNF414, TCF3, ZNF470, KLF1 |
| GOTERM_MF_DIRECT | GO:0003676~nucleic acid binding | 8 | 4.10 | 0.82186083 | AGGF1, ZSCAN30, ZNF223, WRN, ZNF221, ZFP28, KLF1, ZNF470 |
| UP_SEQ_FEATURE | zinc finger region:C2H2-type 10 | 3 | 1.54 | 0.83851486 | ZNF221, ZFP28, ZNF470 |
| UP_SEQ_FEATURE | domain:KRAB | 3 | 1.54 | 0.84090051 | ZNF223, ZNF221, ZNF470 |
| UP_SEQ_FEATURE | zinc finger region:C2H2-type 6 | 4 | 2.05 | 0.85916926 | ZSCAN30, ZNF221, ZFP28, ZNF470 |
| GOTERM_BP_DIRECT | GO:0006351~transcription, DNA-templated | 16 | 8.21 | 0.86594334 | PTOV1, KLF17, ZNF221, ZFP28, TET1, MYCN, HOXD9, HIF1AN, ZSCAN30, ZNF223, ATXN7, ZNF414, TNIP2, TCF3, ZNF470, KLF1 |
| UP_KEYWORDS | Transcription regulation | 18 | 9.23 | 0.89746565 | PTOV1, TAF5, KLF17, ZNF221, ZFP28, TET1, MYCN, HOXD9, HIF1AN, ZSCAN30, ZNF223, ATXN7, IRF3, ZNF414, TNIP2, TCF3, ZNF470, KLF1 |
| UP_KEYWORDS | Transcription | 18 | 9.23 | 0.91962457 | PTOV1, TAF5, KLF17, ZNF221, ZFP28, TET1, MYCN, HOXD9, HIF1AN, ZSCAN30, ZNF223, ATXN7, IRF3, ZNF414, TNIP2, TCF3, ZNF470, KLF1 |
| GOTERM_MF_DIRECT | GO:0003677~DNA binding | 12 | 6.15 | 0.92255128 | HJURP, ZNF223, ZIC5, IRF3, ZNF414, WRN, ZNF221, ZFP28, TET2, ZNF470, TCF3, MYCN |
| UP_KEYWORDS | Nucleus | 42 | 21.54 | 0.93271302 | PTOV1, AKAP13, ANKLE1, CUL3, METTL22, HIF1AN, HJURP, ZNF223, AHNAK2, USP36, MTMR6, NRG1, TNIP2, ZNF470, TCF3, CHUK, GEMIN4, NUP58, NLRP6, TAF5, FLT4, MICAL3, KLF17, ZNF221, WRN, ZFP28, TET1, MYCN, HOXD9, IGFN1, SYNE2, ZSCAN30, CDK11A, ATXN7, ZIC5, CAPG, RBMXL1, AKAP6, IRF3, CDK11B, ZNF414, KLF1 |
| **Annotation Cluster 25** | **Enrichment Score: 0.2132** | | | | |
| **Category** | **Term** | **Count** | **%** | ***p* value** | **Genes** |
| INTERPRO | IPR020472:G-protein beta WD-40 repeat | 3 | 1.54 | 0.22612056 | CDRT1, TAF5, SPAG16 |
| INTERPRO | IPR019775:WD40 repeat, conserved site | 3 | 1.54 | 0.47867268 | CDRT1, TAF5, SPAG16 |
| INTERPRO | IPR017986:WD40-repeat-containing domain | 4 | 2.05 | 0.56649261 | CDRT1, TAF5, VPS11, SPAG16 |
| INTERPRO | IPR015943:WD40/YVTN repeat-like-containing domain | 4 | 2.05 | 0.62196994 | CDRT1, TAF5, VPS11, SPAG16 |
| UP_SEQ_FEATURE | repeat:WD 4 | 3 | 1.54 | 0.70093103 | CDRT1, TAF5, SPAG16 |
| UP_SEQ_FEATURE | repeat:WD 3 | 3 | 1.54 | 0.72852059 | CDRT1, TAF5, SPAG16 |
| INTERPRO | IPR001680:WD40 repeat | 3 | 1.54 | 0.73340629 | CDRT1, TAF5, SPAG16 |
| UP_SEQ_FEATURE | repeat:WD 1 | 3 | 1.54 | 0.7451013 | CDRT1, TAF5, SPAG16 |
| UP_SEQ_FEATURE | repeat:WD 2 | 3 | 1.54 | 0.7451013 | CDRT1, TAF5, SPAG16 |
| UP_KEYWORDS | WD repeat | 3 | 1.54 | 0.74546569 | CDRT1, TAF5, SPAG16 |
| SMART | SM00320:WD40 | 3 | 1.54 | 0.76323488 | CDRT1, TAF5, SPAG16 |
| Annotation Cluster 26 | Enrichment Score: 0.2088 | | | | |
| **Category** | **Term** | **Count** | **%** | ***p* value** | **Genes** |
| UP_KEYWORDS | SH3 domain | 3 | 1.54 | 0.60388459 | DOCK1, ARHGEF37, TRIP10 |
| INTERPRO | IPR001452:Src homology-3 domain | 3 | 1.54 | 0.62408876 | DOCK1, ARHGEF37, TRIP10 |
| SMART | SM00326:SH3 | 3 | 1.54 | 0.62727792 | DOCK1, ARHGEF37, TRIP10 |
| **Annotation Cluster 27** | **Enrichment Score: 0.1692** | | | | |
| **Category** | **Term** | **Count** | **%** | ***p* value** | **Genes** |
| BIOCARTA | h_hivnefPathway:HIV-I Nef: negative effector of Fas and TNF | 3 | 1.54 | 0.144481 | CDK11A, CDK11B, CHUK |
| GOTERM_BP_DIRECT | GO:0006468~protein phosphorylation | 6 | 3.08 | 0.45536692 | P2RX7, CDK11A, AKAP13, CDK11B, CHUK, AATK |
| UP_KEYWORDS | Kinase | 8 | 4.10 | 0.54273496 | CDK11A, FLT4, AKAP6, AKAP13, HGS, CDK11B, CHUK, AATK |
| GOTERM_MF_DIRECT | GO:0004672~protein kinase activity | 4 | 2.05 | 0.65661422 | CDK11A, CDK11B, CHUK, AATK |
| UP_SEQ_FEATURE | domain:Protein kinase | 5 | 2.56 | 0.67048886 | CDK11A, FLT4, CDK11B, CHUK, AATK |
| INTERPRO | IPR000719:Protein kinase, catalytic domain | 5 | 2.56 | 0.69469928 | CDK11A, FLT4, CDK11B, CHUK, AATK |
| UP_KEYWORDS | Serine/threonine-protein kinase | 4 | 2.05 | 0.71945418 | CDK11A, CDK11B, CHUK, AATK |
| UP_SEQ_FEATURE | nucleotide phosphate-binding region:ATP | 9 | 4.62 | 0.73525591 | NLRP6, CDK11A, FLT4, DYNC2H1, CDK11B, ATP10D, WRN, CHUK, AATK |
| INTERPRO | IPR011009:Protein kinase-like domain | 5 | 2.56 | 0.75708265 | CDK11A, FLT4, CDK11B, CHUK, AATK |
| UP_SEQ_FEATURE | active site:Proton acceptor | 6 | 3.08 | 0.77069757 | CDK11A, FLT4, DHRS4L2, CDK11B, CHUK, AATK |
| UP_SEQ_FEATURE | binding site:ATP | 5 | 2.56 | 0.7814549 | CDK11A, FLT4, CDK11B, CHUK, AATK |
| INTERPRO | IPR008271:Serine/threonine-protein kinase, active site | 3 | 1.54 | 0.8061796 | CDK11A, CDK11B, CHUK |
| GOTERM_MF_DIRECT | GO:0005524~ATP binding | 12 | 6.15 | 0.83724387 | NLRP6, P2RX7, TTLL10, CCT6B, CDK11A, FLT4, DYNC2H1, CDK11B, ATP10D, WRN, CHUK, AATK |
| UP_KEYWORDS | ATP-binding | 11 | 5.64 | 0.85176126 | NLRP6, TTLL10, CCT6B, CDK11A, FLT4, DYNC2H1, CDK11B, ATP10D, WRN, CHUK, AATK |
| UP_KEYWORDS | Nucleotide-binding | 14 | 7.18 | 0.87006665 | NLRP6, FLT4, ATP10D, ARL16, WRN, TTLL10, CCT6B, CDK11A, DYNC2H1, CDK11B, TUBA3E, VPS11, CHUK, AATK |
| GOTERM_MF_DIRECT | GO:0004674~protein serine/threonine kinase activity | 3 | 1.54 | 0.87028809 | CDK11A, CDK11B, AATK |
| INTERPRO | IPR017441:Protein kinase, ATP binding site | 3 | 1.54 | 0.88522468 | FLT4, CHUK, AATK |
| SMART | SM00220:S_TKc | 3 | 1.54 | 0.88738342 | CDK11A, CDK11B, CHUK |
| **Annotation Cluster 28** | **Enrichment Score: 0.1639** | | | | |
| **Category** | **Term** | **Count** | **%** | ***p* value** | **Genes** |
| INTERPRO | IPR013083:Zinc finger, RING/FYVE/PHD-type | 5 | 2.56 | 0.63928797 | JADE2, HGS, VPS11, RNF213, TRAF3 |
| UP_SEQ_FEATURE | zinc finger region:RING-type | 3 | 1.54 | 0.64595732 | VPS11, RNF213, TRAF3 |
| INTERPRO | IPR001841:Zinc finger, RING-type | 3 | 1.54 | 0.78042372 | VPS11, RNF213, TRAF3 |
| **Annotation Cluster 29** | **Enrichment Score: 0.1337** | | | | |
| **Category** | **Term** | **Count** | **%** | ***p* value** | **Genes** |
| GOTERM_CC_DIRECT | GO:0043025~neuronal cell body | 4 | 2.05 | 0.59696937 | P2RX7, APOB, GRIA1, LRP4 |
| UP_KEYWORDS | Palmitate | 3 | 1.54 | 0.81251688 | P2RX7, APOB, GRIA1 |
| UP_KEYWORDS | Lipoprotein | 7 | 3.59 | 0.81865814 | P2RX7, APOB, GRIA1, RAET1L, GHRL, INPP5B, LRP4 |
| **Annotation Cluster 30** | **Enrichment Score: 0.0564** | | | | |
| **Category** | **Term** | **Count** | **%** | ***p* value** | **Genes** |
| GOTERM_MF_DIRECT | GO:0004842~ubiquitin-protein transferase activity | 3 | 1.54 | 0.81665011 | CUL3, RNF213, TRAF3 |
| GOTERM_BP_DIRECT | GO:0016567~protein ubiquitination | 3 | 1.54 | 0.86662918 | CUL3, RNF213, TRAF3 |
| UP_KEYWORDS | Ubl conjugation pathway | 4 | 2.05 | 0.95714435 | CUL3, USP36, RNF213, TRAF3 |
