## Supplementary material for "Whole exome sequencing identifies novel DYT1 dystonia-associated genome variants as potential disease modifiers": Figure S1

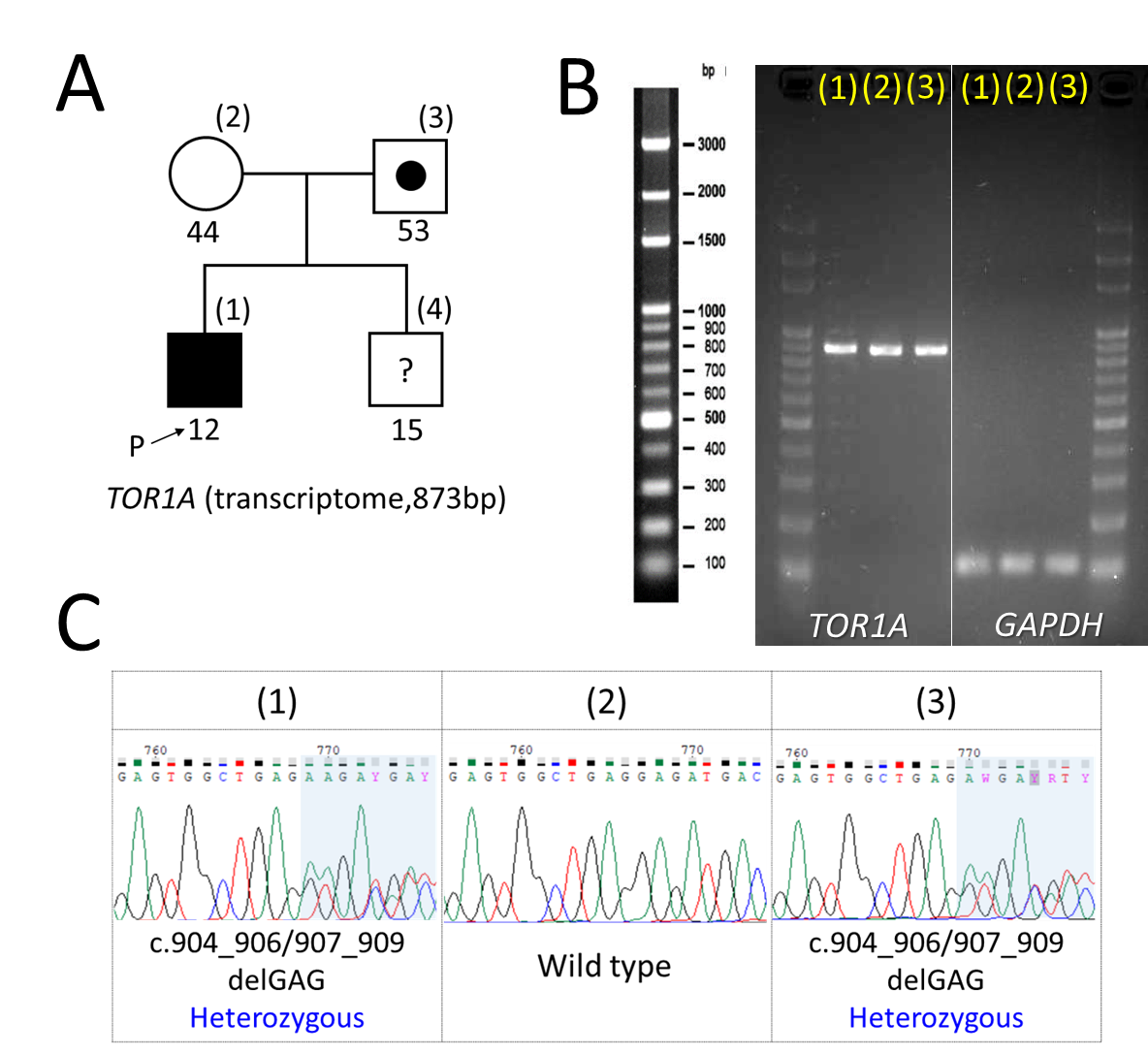

**Figure S1. First patient and the core family pedigrees (A), cDNA product of TOR1A gene (transcriptome, 873bp) and GAPDH gene (internal control, 110bp) (B) and Sanger sequencing data (C) of TOR1A gene (transcriptome, 873 bp).** (A) The arrow points out the first proband. The numbers within parentheses are the order of Sanger sequencing data and the numbers under the box/circle show the age (years old). The question marks within the box indicate the unknown status because we don’t have the cDNA sample for study.
